# Decoding antibiotic modes of action from multimodal cellular responses

**DOI:** 10.64898/2026.03.31.715570

**Authors:** Joshua Hesse, Dominik Schum, Laura Leidel, Leonard R. Gareis, Jennifer Herrmann, Rolf Müller, Stephan A. Sieber

## Abstract

Antibiotic resistance continues to rise, yet most new drug candidates act through long-established targets. Faster mode of action (MoA) assessment would enable more effective prioritization of screening hits and help identify compounds with novel mechanisms. In this study, we aimed to develop a scalable framework for MoA inference from antibiotic-induced cellular response profiles in *Escherichia coli*. We generated a multimodal dataset spanning more than 50 antibiotics, including proteome profiles, chemical structure descriptors, inhibitory concentrations and growth dynamics, and used it to build MAPPER (Mode of Action Prediction via Proteomics-Enhanced Representation), a framework comprising a fixed multimodal predictor and an uncertainty module. MAPPER accurately classified antibiotics across nine mechanistic classes, flagged compounds with likely novel mechanisms and retained predictive power in proteomics-only transfer experiments across mass spectrometry platforms and external data. Together, these results establish MAPPER as an innovative tool for MoA prediction and novelty detection, enabling prioritization of antibacterial candidates with distinct mechanisms.

## Introduction

Antimicrobial resistance continues to rise while the approval rate of new antibiotics steadily declines [1, 2]. Antibiotic development pipelines remain focused on derivatives of established scaffolds, prioritizing compounds that act through familiar targets rather than expanding the diversity of mechanisms available for therapy [3, 4]. This limited exploration of the mechanistic space contributes to cross-resistance and reduces the therapeutic effects of new antibiotics, while resistance mechanisms continue to develop [4–7]. Gram-negative bacteria pose an additional challenge for treatment, combining restricted permeability with an extensive repertoire of resistance pathways [8], resulting in a widening gap between clinical need and discovery pipelines [9].

A major bottleneck in the antibiotic discovery pipeline is the determination of an antibiotic’s mode of action (MoA)[10–12]. Upon discovery of a new antibiotic scaffold, target identification often relies on experimentally demanding methods such as sequencing of resistant strains [13, 14], activity-based protein profiling (ABPP) [15, 16], or chemical–genetic interaction profiles (CGIPs) [17], followed by resource-intensive target confirmation experiments [11, 18]. This investment is particularly costly when the elucidated MoA ultimately corresponds to an already known mechanism despite a structurally novel compound class. Faster and more scalable MoA elucidation would therefore enable broader chemical exploration and help prioritize candidates with genuinely novel activities, which are urgently needed to counter the rising antimicrobial resistance crisis [16].

In this context, machine learning (ML) offers a scalable route to MoA assessment by extracting distributed mechanistic signals from high-dimensional cellular response profiles. This is particularly valuable where informative responses are subtle, multimodal, and difficult to interpret through manual or pathway-by-pathway analysis alone.

ML has begun to address this challenge by leveraging high-dimensional cellular readouts for MoA inference [19, 20]. Recent work has shown that antibiotic MoAs can be inferred from transcriptomic signatures [21], proteome-wide response profiles [22, 23], metabolomic perturbation data [24–26], and image-based bacterial phenotypes [27]. Each modality, however, has limitations. Transcriptomic signatures can be indirect and may miss post-transcriptional or protein-level adaptation [28, 29]. Metabolomics is informative only when antibiotic treatment produces measurable metabolic perturbations [24, 25]. Imaging captures rich phenotypes but often lacks direct mechanistic interpretability and is particularly challenging in bacteria, where morphological responses are comparatively constrained and can overlap across MoAs [30–32]. Proteomics provides a more direct view of protein-level responses and has shown promise for MoA classification [22, 23, 33], but prior proteomics-based studies have often relied on small reference panels or reduced marker sets, limiting scalability and integration with complementary phenotypic context.

Together, these considerations motivate the development of integrative approaches that combine the mechanistic specificity of proteomics with complementary chemical and phenotypic information, while remaining scalable and applicable to compounds with uncertain or novel MoAs. In the following, we present MAPPER, a multimodal framework for antibiotic MoA prediction in the Gram-negative strain *Escherichia coli* (*E. coli*) that integrates proteome-level responses with auxiliary experimental and chemical descriptors, and we assess its performance and robustness across multiple evaluation settings.

## Results

### MAPPER framework overview

We established MAPPER (Mode of Action Prediction via Proteomics-Enhanced Representation), a multimodal framework for antibiotic MoA inference in *E. coli* (Figure 1). In the prediction branch (Figure 1a), compound-specific phenotypic, proteomic, and structural data are combined into a multimodal fingerprint and paired with text embeddings of candidate MoA descriptions, reformulating MoA prediction as a description–feature matching task suited to limited sample size. In the second branch (Figure 1b), metadata derived from these predictions is used by an uncertainty module to flag compounds whose responses deviate from the mechanistic space represented in the training data. The framework was developed on a standardized reference dataset spanning nine antibiotic classes (Figure 1c), which provides the basis for both classification and novelty-aware decision support.

**Fig. 1.**
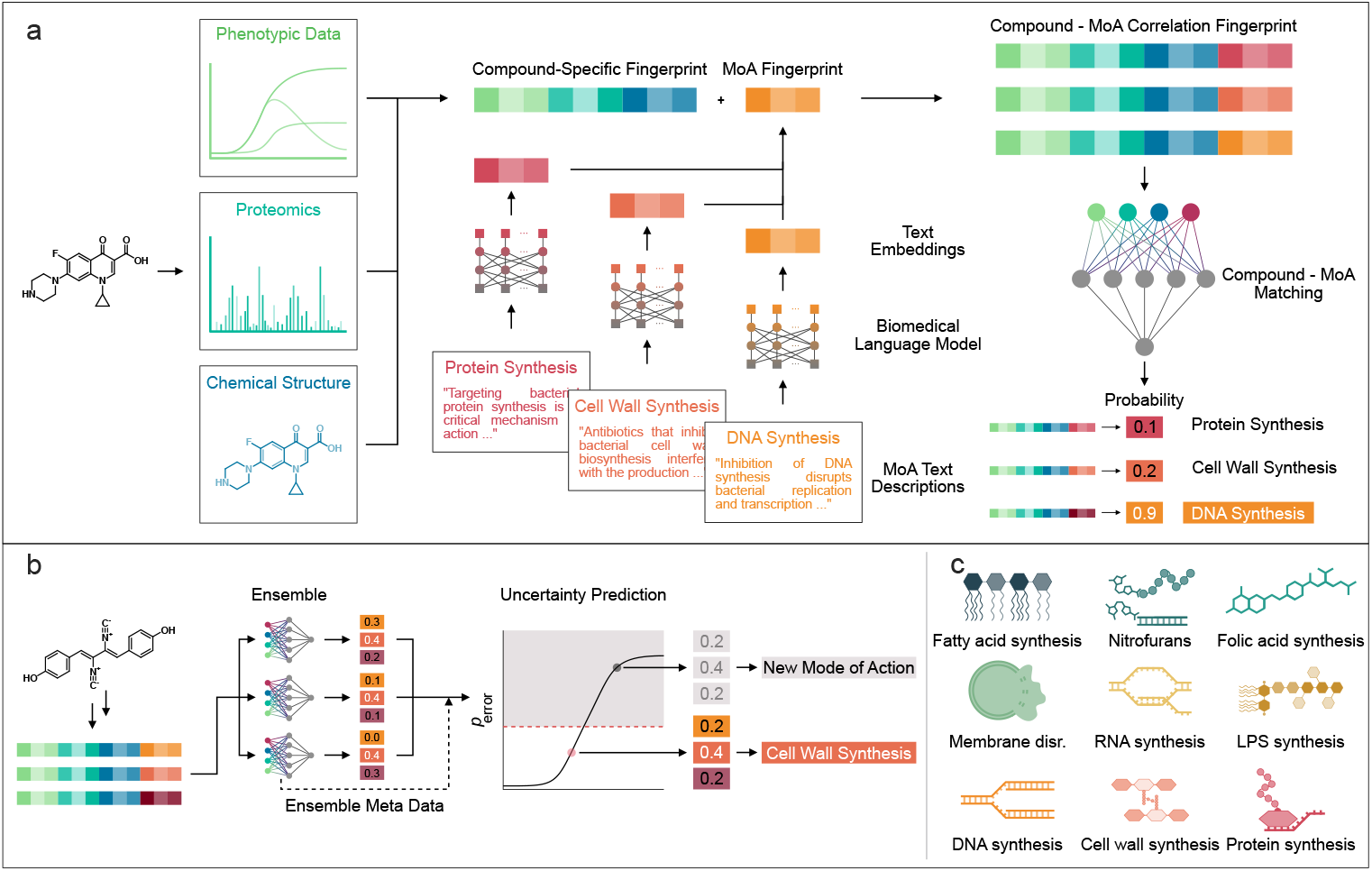
MAPPER framework overview: (a) Phenotypic, proteomic, and structural data is generated for each antibiotic compound and combined to a compound-specific fingerprint. Text descriptions of all MoAs are converted to text embeddings and combined with the compound fingerprints. The combined fingerprints are used as input for a machine learning model that predicts the probability of the feature- and text-based fingerprints to match. (b) Metadata of the prediction is used to predict the model’s uncertainty, identifying compounds with potentially novel MoAs during inference. (c) The library comprises 51 compounds with nine antibiotic MoAs: fatty acid synthesis inhibition, nitrofuran antibiotics, folic acid synthesis inhibition, membrane disruption, RNA synthesis inhibition, lipopolysaccharide synthesis inhibition, DNA synthesis inhibition, cell wall synthesis inhibition, and protein synthesis inhibition.

### Generation of antibiotic profile dataset

We assembled a library of 68 antibiotics with reported activity against *E. coli*, encompassing nine mechanistic classes and several structurally diverse scaffolds. Compounds lacking measurable antibacterial activity across tested concentrations or exhibiting strong inoculum effects were excluded from downstream proteomic analysis. The final model-development dataset consisted of 51 fully characterized antibiotics. Vendor sources and the full compound library, including its use in model development or subsequent validation, are listed in Supplementary Tables 1 and 2.

For all compounds with detectable activity, minimum inhibitory concentrations (MICs) were determined and used to guide the design of growth inhibition assays at higher cell densities (Figure 2a). Growth curves revealed class-dependent differences in inhibition kinetics for several groups (Figure 2b), while other classes showed more heterogeneous responses (Ext. Figure 1). These measurements, together with MICs, were included as quantitative phenotypic features.

**Fig. 2.**
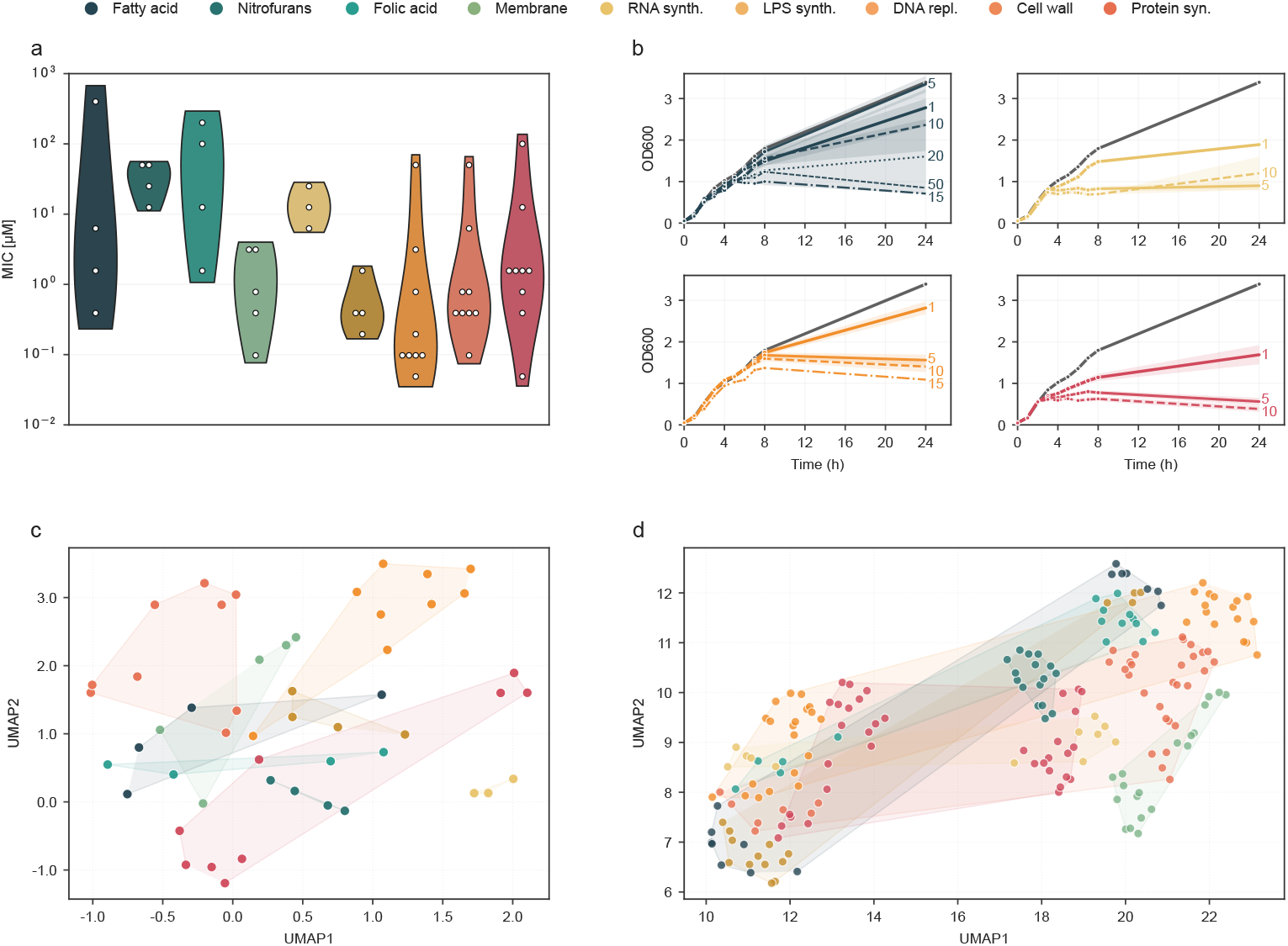
Biochemical dataset generation: (a) Distribution of MIC values for each MoA class. (b) Growth curve effects of DNA replication, fatty acid synthesis, protein synthesis, and RNA synthesis inhibitor classes. Growth controls are shown in gray. Each data series shows the average values over all members of the respective class with the given MIC factor. MIC factors are 1x, 5x, 10x, 15x, 20x, 50x MIC. (c) Unsupervised UMAP of the ECFP fingerprints of all compounds, with shaded areas representing the structural space of the different classes. (d) Unsupervised UMAP of the full proteome fingerprints for all compounds and all replicates, with shaded areas representing the proteomic space of the different classes.

Chemical structure information was captured using Extended-Connectivity Fingerprints (ECFPs). A UMAP projection of the fingerprints (Figure 2c) showed compact clustering for some classes and broader distributions for others, reflecting differences in structural diversity within and across MoA groups.

To characterize proteome-level effects, full proteome response data for all active compounds was generated by quantitative mass spectrometry using standardized bottom-up sample processing and acquisition workflows. Samples were prepared by two different operators to ensure that the dataset captured realistic technical variability rather than relying on a single-operator workflow. An unsupervised UMAP embedding of all biological replicates (Figure 2d) revealed a clear operator-associated separation. Importantly, because most MoA classes contained samples originating from both operators, any operator-dependent variation was not systematically aligned with the labels, meaning that downstream models should learn to discount operator effects rather than rely on them. Thus, the dataset remained well suited for MoA modeling while explicitly incorporating technical variation.

In addition to these bacterial features, we generated orthogonal morphological– transcriptional proxy embeddings for each compound using the InfoAlign model [34], which projects chemical structures into a pretrained joint embedding space learned from large-scale mammalian perturbation data.

Taken together, these measurements yield a structurally and mechanistically diverse reference panel with matched phenotypic and proteomic readouts, providing a strong basis for training and objectively evaluating MoA prediction models.

### Text augmentation for dataset inflation

The model-development dataset comprises 51 compounds with four biological replicates each, yielding only about 200 samples, leaving the number of observations small relative to the dimensionality of the feature space and thus making learning statistically difficult[35]. We therefore reformulated MoA prediction as a binary description–feature matching task, pairing each biochemical fingerprint with each of the nine correct or incorrect candidate MoA descriptions (Figure 3a)[36, 37], which expanded the dataset to roughly 1,800 data points. To further increase diversity, we generated ten augmented variants of each MoA description (Figure 3b)[38], yielding roughly 18,000 rows in total.

**Fig. 3.**
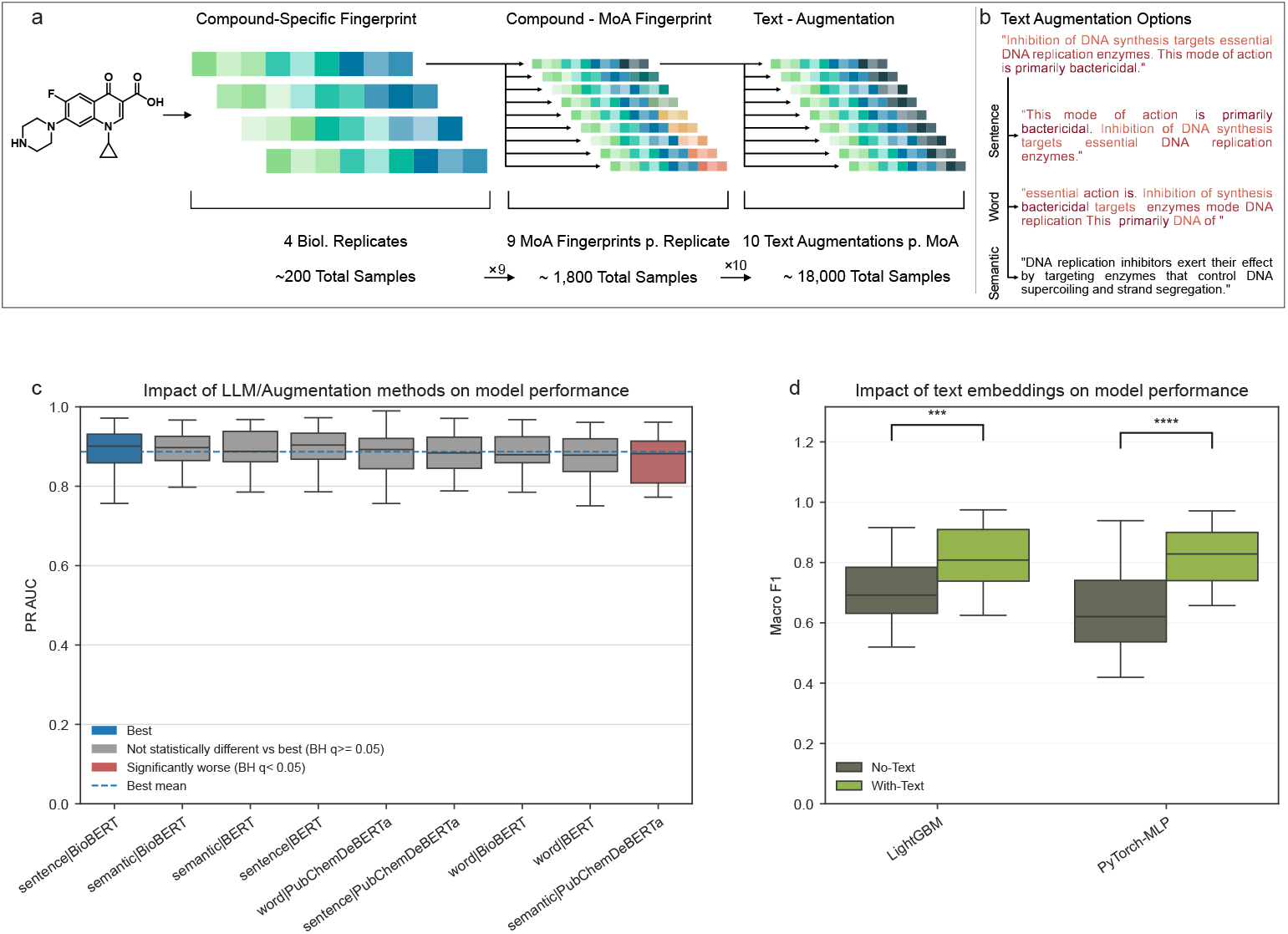
Text-based fingerprints: (a) Four biological replicate fingerprints were generated for each compound. Each biological fingerprint was combined with nine MoA description fingerprints. Each combined fingerprint was then augmented ten times by using slightly changed representations of the same text descriptions. (b) Three types of text augmentations were tested. Sentence-based augmentation shuffles the sentences of a given text description ten times. Word-based augmentation shuffles all words randomly for a given description. For semantic augmentation, the text was rewritten ten times. (c) PR AUC across LLM/augmentation methods (PyTorch-MLP, Text Embedding + Proteomics). Boxplots show median (center line), IQR (box), whiskers (1.5×IQR) and outliers (points). Overall differences across methods were assessed by repeated-measures ANOVA per metric (*n* = 25 cross-validation cycles), with Bonferroni correction across PR AUC, ROC AUC, and MCC (*α*_Bonf_ = 0.0167). Omnibus *p*-value for PR AUC was = 1.96 *×* 10^*−*3^. Tukey HSD pairwise comparisons were then used for PR AUC coloring (best = blue; not different = gray; worse = red). (d) Macro F1 for proteomics-only models With-Text vs No-Text, grouped by model (LightGBM, PyTorch-MLP). Boxplots as in (c). The overall four-group comparison (model *×* text condition) was assessed by Friedman test (*p* = 1.04 *×* 10^*−*6^; *n* = 25 balanced cycles). Brackets denote within-model paired *t* -tests comparing With-Text vs No-Text; stars indicate Bonferroni-adjusted significance.

We evaluated three BERT-family language models (Bidirectional Encoder Representations from Transformers (BERT), BioBERT, PubChemDeBERTa) as text embedding encoders for the MoA descriptions and three augmentation strategies (sentence shuffling, word shuffling, or semantic rewriting) in a 5 × 5 cross-validation scheme using a multilayer perceptron (MLP) as the prediction model [39–41]. Model choice and augmentation type had minimal effects on performance (Figure 3c), and exact ranking changes based on the underlying ML model and performance metric used (Supplementary Figure 1). Because semantic augmentation offers greater diversity than sentence shuffling while preserving readability better than word shuffling, we chose semantic augmentation in combination with BioBERT for all subsequent experiments.

To assess whether augmentation improved predictive performance, we compared models trained on the original 200 biological samples with those trained on the augmented dataset. Both LightGBM and a custom neural network achieved substantially higher performance using the augmented data (Figure 3d), supporting its use in subsequent analyses. This demonstrates that task reformulation in combination with text-based augmentation effectively compensates for limited experimental sample size and unlocks the full predictive capacity of more training-size-dependent model classes.

### Model and feature set selection

We next evaluated how different feature combinations contributed to MoA prediction using 5 × 5 cross-validation across MIC, growth curve, proteomics, chemical fingerprint, and InfoAlign features. Predictive performance increased with feature complexity, with proteomic, structural, and InfoAlign features each providing substantial gains over MIC or growth curve data alone (Figure 4a). Across random forests, gradient boosting methods, neural networks, and TabM[42], the best-performing feature sets consistently included structural, proteomic, and InfoAlign descriptors (Supplementary Table 7).

**Fig. 4.**
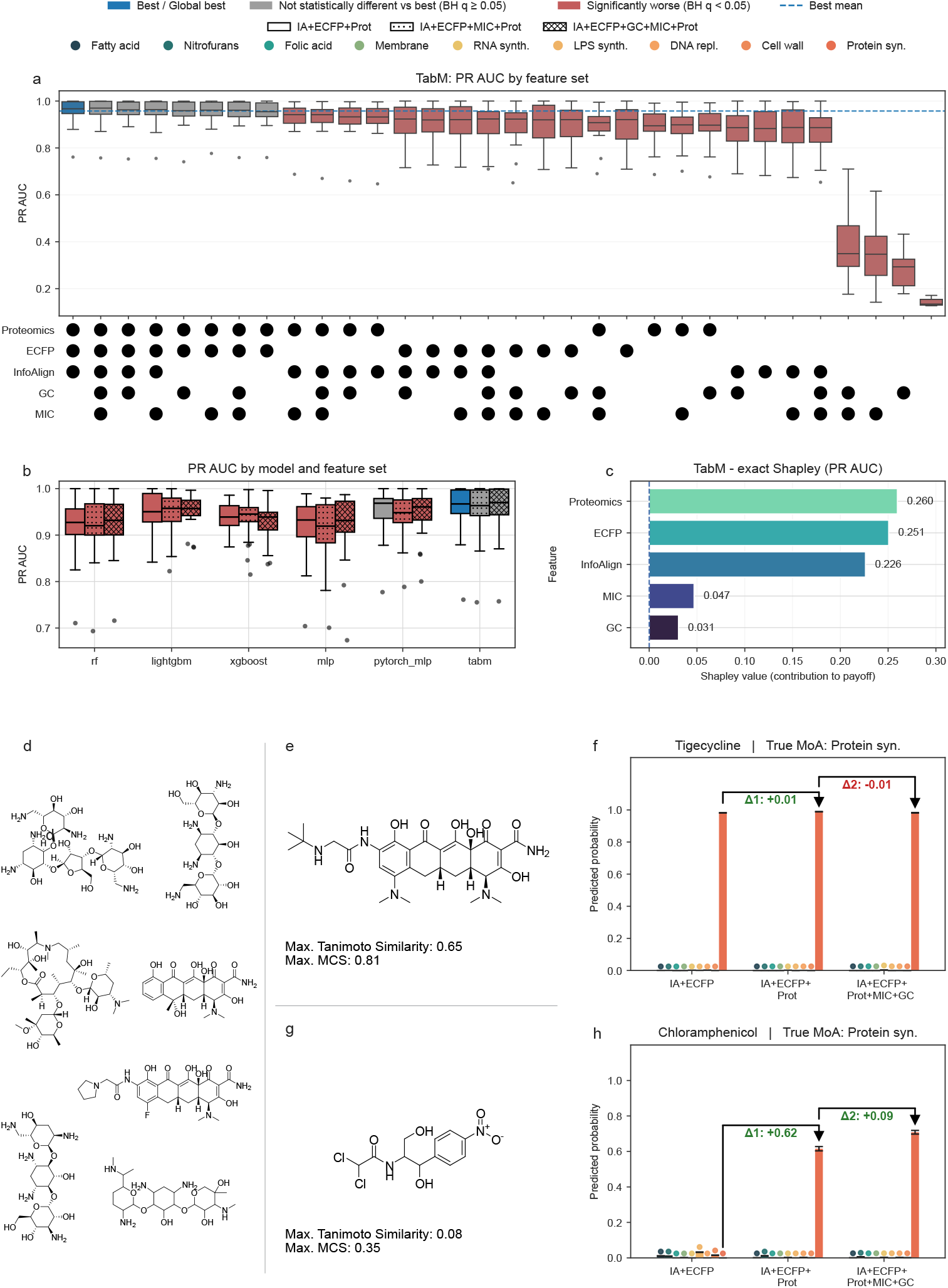
Feature-dependent mode of action prediction: (a) TabM PR AUC across feature sets (Text Embedding always included). Boxplots show median (center line), IQR (box), whiskers (1.5×IQR) and outliers (points). Feature inclusion is shown by the dot matrix (Proteomics, ECFP, InfoAlign, GC, MIC). Friedman test across feature sets (*n* = 25 cross-validation cycles): *p* = 1.84 *×* 10^*−*73^. Post-hoc Conover–Friedman pairwise comparisons with Benjamini–Hochberg correction (*q <* 0.05) determine coloring vs the best feature set (blue = best; grey = not different; red = worse). (b) PR AUC across six model types for three top feature sets. Boxplots as in (a). Friedman test across all model × feature-set conditions (18 conditions; *n* = 25 balanced cross-validation cycles): *p* = 5.61 10^*−*23^. Post-hoc Conover–Friedman pairwise comparisons with Benjamini–Hochberg correction (*q <* 0.05) determine coloring vs the global best. Feature-set patterns are indicated by boxed symbols in the legend. (c) Exact Shapley values for TabM PR AUC computed from all 32 feature coalitions (Proteomics, ECFP, CellPaint, MIC, GC); bars show contribution to model performance. Chemical structures of the remaining protein synthesis inhibitors. (e, g) Structure of tigecycline & chloramphenicol, maximum Tanimoto similarity and maximum common substructure score compared to the remaining class members. (f, h) Tigecycline and chloramphenicol predicted MoA probabilities using InfoAlign and chemical structure fingerprints vs adding proteomics vs all feature modalities. Bars show mean ± SEM across *n* = 200 replicate predictions per MoA; colored dots denote MoA categories. Arrows and labels indicate the delta in mean predicted probability for the true MoA due to addition of proteomics (Δ1) and MIC & GC data (Δ2).

Comparison of the six model classes on the three strongest feature sets identified TabM as the best overall model, with all three top feature sets outperforming most alternative model–feature combinations (Figure 4b, Supplementary Figure 2). Exact Shapley values computed for TabM confirmed proteomics as the strongest additive contributor, followed by structural and InfoAlign features, whereas MIC and growth curve features had comparatively modest effects (Figure 4c).

To examine how biological data aids predictions for structurally distinct compounds, we compared leave-one-out (LOO) predictions with and without biological features. For compounds with high structural similarity to their class, such as tige-cycline among the protein synthesis inhibitors, predictions were accurate as long as fingerprints based on structural information were present (Figure 4d-f). For structurally distinct protein synthesis inhibitors such as chloramphenicol (Figure 4g-h), proteomics was required for correct classification. No prediction was possible when proteomics data was absent. Addition of MIC and growth curve data further increased correct prediction probability. The same effect was observed for structurally distinct inhibitors of other classes, such as gepotidacin, a structurally distinct DNA synthesis inhibitor, and Debio-1452-NH_2_, a fatty acid synthesis inhibitor (Ext. Figure 2). Therefore, we decided to use the full feature set whenever possible for subsequent applications, as biological data positively affects predictions of structurally novel molecules. Based on the combined results of Figures 3 and 4, we fixed a semantic+BioBERT TabM classifier using the full multimodal feature set as the base predictor used in MAPPER.

To test whether classical pathway enrichment alone would suffice for MoA inference, we performed STRING-based analysis of significantly upregulated proteins for one representative compound from each of nine mechanistic classes (Ext. Figure 3)[43]. We chose this comparison because pathway enrichment is a widely used and biologically interpretable first-pass strategy for linking proteomic changes to perturbed cellular processes. Enriched biological processes were detected for five compounds, but only DNA synthesis and protein synthesis inhibitors yielded enrichments aligned with the expected MoA; four compounds showed no significant enrichment (Ext. Figure 4). In contrast, the multimodal model correctly classified all nine compounds (Ext. Figure 3, 4). Thus, although enrichment can be informative in specific cases, it is not sufficiently sensitive or general for MoA determination, whereas ML can exploit subtler distributed proteomic signals.

### Uncertainty Estimation for Novel MoA Detection

A central goal of this study was not only to assign compounds to known mechanistic classes, but also to recognize when a compound falls outside the mechanistic space represented in the training data. Because the fixed MAPPER predictor is trained only on known MoAs, it will still return one of these classes at inference time, even for compounds with genuinely novel mechanisms. We therefore evaluated uncertainty estimation strategies that identify when such assignments should be treated as unreliable (Figure 5). For benchmarking, we used two complementary settings: LOO, which approximates in-distribution inference on unseen compounds from known classes, and leave-one-class-out (LOCO), which simulates mechanistic novelty by excluding an entire MoA class from training. Correct LOO predictions were used as examples of reliable in-distribution predictions, whereas LOCO predictions were used as examples of novelty-like predictions for which the model should be uncertain.

**Fig. 5.**
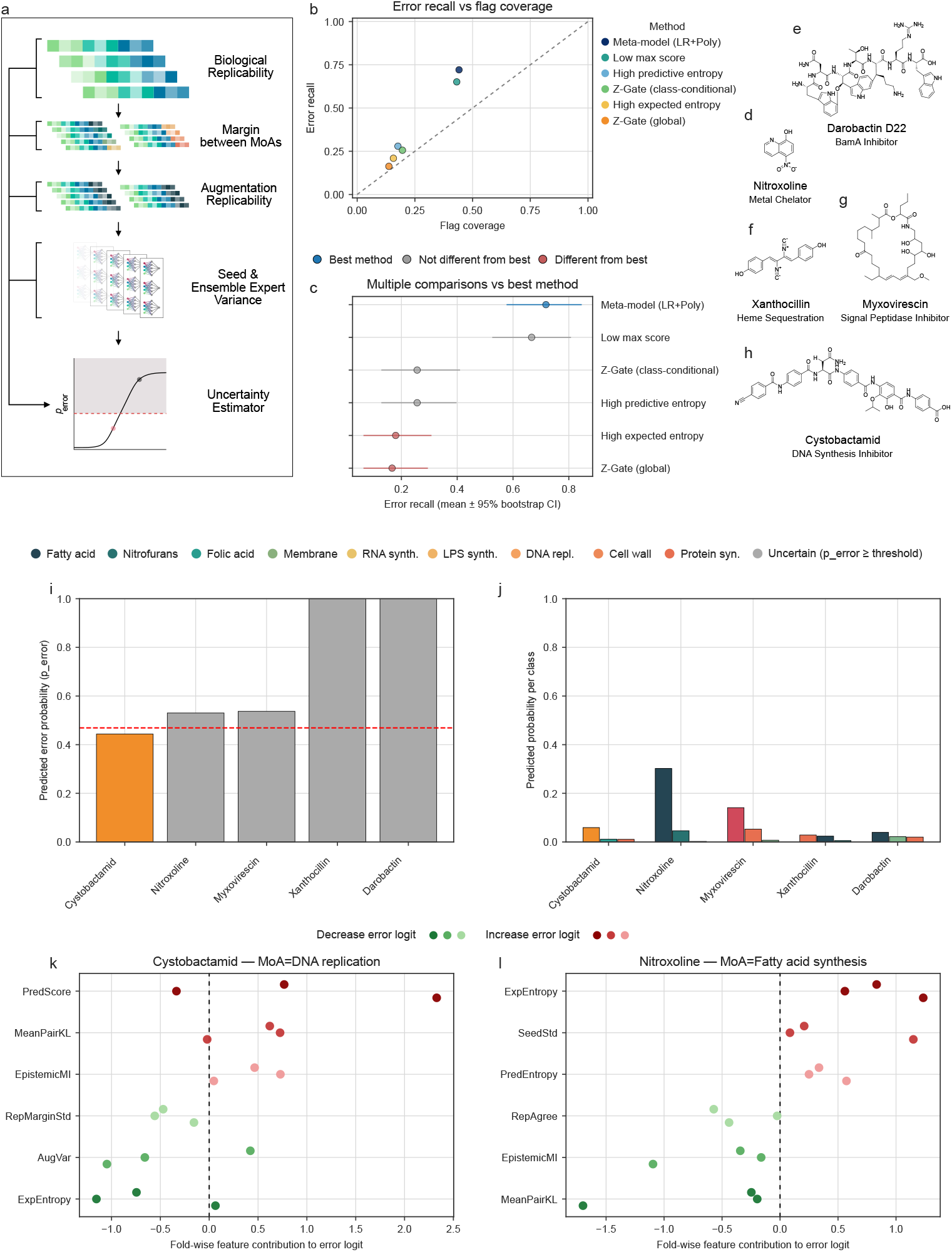
Uncertainty estimation: (a) Uncertainty-estimation workflow. The fixed MAPPER predictor first produces class scores for each compound. From these outputs we derive uncertainty features describing top-class support and class separation, entropy of the class distribution, disagreement across perturbation-level predictions, and consistency across biological replicates or other perturbations. These features are combined into an uncertainty score, above which a sample is flagged as uncertain. (b) Error recall vs flag coverage for the six uncertainty methods benchmarked on the LOO/LOCO setting: low max score, high predictive entropy, high expected entropy, Z-Gate (global), Z-Gate (class-conditional), and the meta-model (LR+Poly). Gray line resembles random picking. (c) Multiple-comparisons plot, showing the best-performing method in blue, methods not significantly different in grey, and significantly worse methods in red. (d-h) Structures and MoAs of held-out antibiotics used for final evaluation. (i) Uncertainty estimates for the held-out antibiotics. Red line indicates the uncertainty threshold, with predictions above threshold deemed uncertain (grey). (j) Top three prediction probabilities for each held-out antibiotic by the MAPPER predictor. (k) and (l) Fold-wise meta-model logit decomposition for cystobactamid (k) and nitroxoline (l). Features shown are the three strongest positive and three strongest negative mean contributors across the calibrated ensemble; red points increase and green points decrease the error logit.

The compared methods included simple score- and entropy-based baselines, disagreement-based scores, two rule-based z-score gates, and a meta-model that combines multiple uncertainty features into a calibrated error probability (Figure 5b–c). These features were derived from the aggregated multiclass output of the MAPPER predictor and its variation across biological replicates, text augmentations, random seeds and ensemble members, capturing score strength and class separation (Pred-Score, Margin), distributional uncertainty, disagreement between perturbation-level predictions, and replicate or perturbation instability. All methods were compared at matched in-distribution conservativeness by fixing the false-positive rate on correct LOO predictions and then measuring how effectively each method flagged erroneous or novelty-like predictions. A logistic regression meta-model achieved the most favorable trade-off between flag coverage and error recall and was therefore combined with the fixed multimodal classifier to define MAPPER.

We then evaluated the full prediction-and-uncertainty framework on a held-out final evaluation set of five antibiotics excluded from all model training, feature selection, and uncertainty-model optimization steps (Figure 5d–h): cystobactamid, a DNA synthesis inhibitor; darobactin D22, which targets outer-membrane protein assembly; myxovirescin, a type II signal peptidase inhibitor; xanthocillin, which acts via dysregulation of heme biosynthesis; and nitroxoline, whose reported mechanism involves metallophore activity [44–49]. These compounds were selected to provide structural novelty relative to the 51-compound model-development set, and four of the five have mechanisms not represented in the training data. Cystobactamid, the only compound with an MoA present in the training set[50–52], was correctly assigned to the DNA synthesis inhibitor class and remained below the uncertainty threshold, consistent with its reported activity (Figure 5i-j). The remaining four compounds were flagged as uncertain by the meta-model, in line with their novel MoAs. Several of these compounds nevertheless showed moderate raw prediction scores, illustrating that class probability alone is not a reliable indicator of prediction confidence.

To understand these differences, we examined the individual uncertainty features contributing to the meta-model. Cystobactamid, the only held-out compound with a mechanism represented in the training set, provides an example of a reliable prediction: although it shows some positive risk contributions from score- and disagreement-related terms, these are outweighed by low expected entropy and replicate-variability signals (AugVar, RepMarginStd), keeping its error probability below threshold (Figure 5k). Nitroxoline, in contrast, represents a novelty-like case: its error probability is driven upward mainly by high expected entropy, seed-to-seed variability, and predictive entropy, while other signals such as replicate agreement, MeanPairKL, and EpistemicMI contribute in the opposite direction, but not strongly enough to bring the predicted error probability below the threshold (Figure 5l). These examples show that the uncertainty model does not simply track prediction score, but integrates score-, entropy-, and stability-related signals to distinguish reliable in-distribution predictions from high-scoring but unstable assignments, thereby helping separate genuinely novel response patterns from those of known classes.

### Prediction on external proteomics datasets

We next tested whether the proteomics branch of MAPPER remains informative under domain shift using a proteomics-only transfer setting. Three antibiotics—kanamycin, imipenem and ciprofloxacin—together with matched controls were re-measured on a different mass spectrometry platform, and the evaluated compounds were excluded from training. We restricted these experiments to proteomics to isolate the contribution of this modality, as a full multimodal model could otherwise exploit structural similarity to compounds already represented in the training set. When applied directly to all protein intensities acquired on the second instrument, the model did not yield confident predictions (Figure 6a). However, restricting both training and inference to proteins showing significant changes relative to controls restored confident and correct MoA assignments for all three antibiotics (Figure 6b). These results indicate that statistically robust proteomic changes preserve the mechanistic signal needed for accurate classification despite platform-specific variation.

**Fig. 6.**
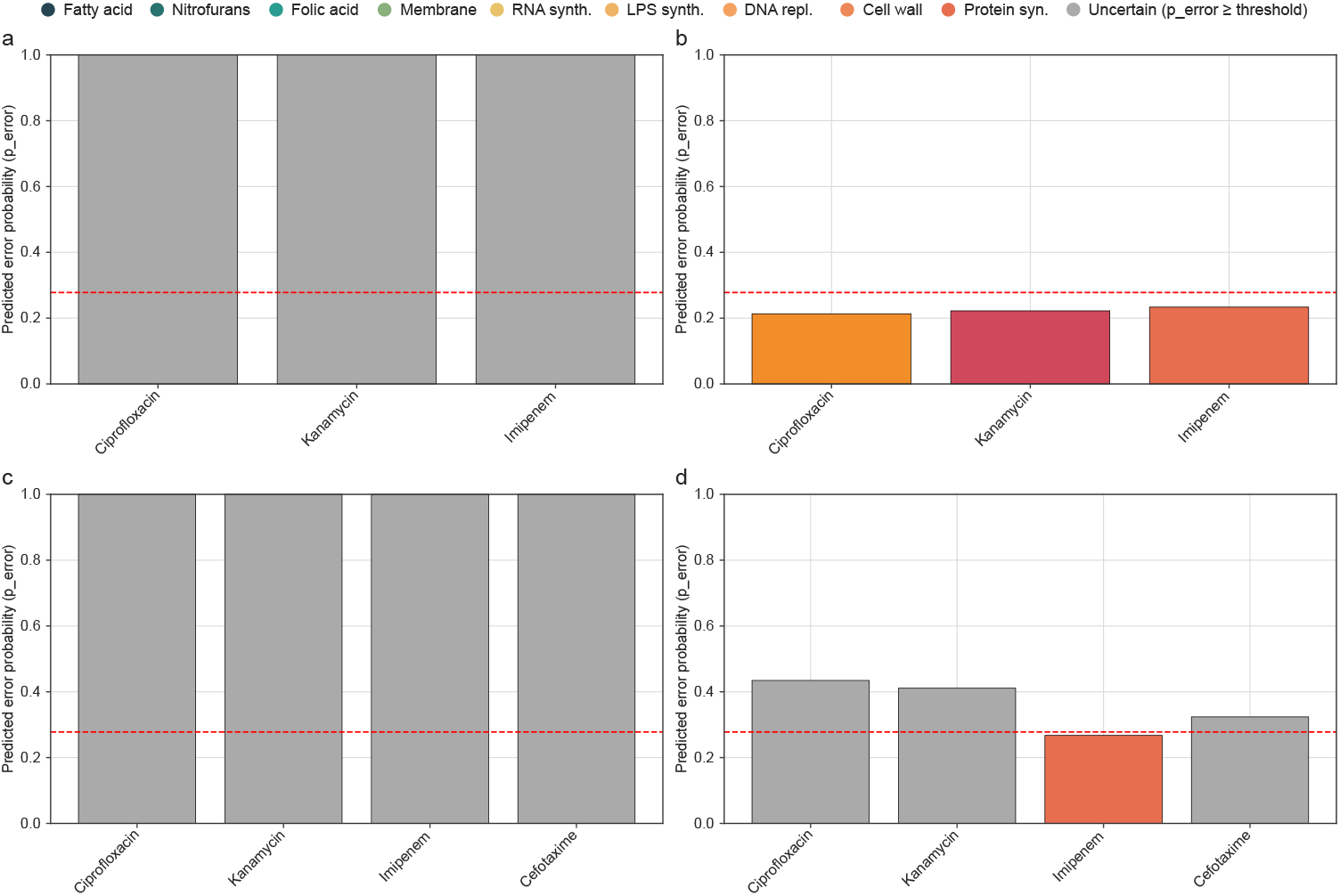
Prediction of external data: (a-b) MoA prediction of held-out antibiotics, using only proteomics features, with the held-out compounds measured on a different mass spectrometer. (a) Prediction and uncertainty estimation when model was used with the unfiltered proteomic fingerprint format. (b) Prediction and uncertainty estimation when only genes with significant changes to controls were used for training and inference. (c-d) MoA prediction of proteomics data for three antibiotics from Subanovic et al. [53], using only proteomics features for training and inference. (c) Prediction and uncertainty estimation when model was used with the unfiltered proteomic fingerprint format. (d) Prediction and uncertainty estimation when only genes with significant changes to controls were used for training and inference. Red line indicates “uncertainty threshold” with predictions above the threshold deemed uncertain (grey).

We next evaluated an external proteomics dataset generated in an independent laboratory. A recent study by Subanovic et al. profiled *E. coli* treated with several antibiotics at subinhibitory concentrations [53], providing a stringent benchmark because it differs from our training data in both laboratory workflow and treatment regime, as our model was trained on inhibitory-concentration responses. When applied directly to all protein abundances from this dataset, the model did not generate confident MoA predictions (Figure 6c), consistent with the cross-instrument experiment. We therefore applied the same significance-filtered framework used above. Under this setting, the model correctly identified the MoA of imipenem while remaining uncertain for the other compounds (Figure 6d), consistent with the external dataset itself, in which imipenem was the only antibiotic inducing a clear proteomic response whereas kanamycin, cefotaxime and ciprofloxacin caused little or no significant change[53].

Together, these experiments demonstrate that restricting analyses to statistically significant protein-level changes enables robust MoA inference even under intentionally unfavorable conditions, including cross-laboratory data and biologically weak perturbations. This highlights our approach’s ability to extract mechanistic signals from proteomics data beyond the specific experimental regime used for training, without hallucinating modes of action where none are detectable.

## Discussion

This study establishes MAPPER (Mode of Action Prediction via Proteomics-Enhanced Representation), a proteomics-anchored multimodal framework for predicting antibiotic MoAs in *E. coli*. MAPPER combines a fixed multimodal predictor with a downstream uncertainty layer to support accurate classification across diverse compound classes while identifying cases in which the data does not justify confident assignment. The resulting dataset, comprising more than 50 antibacterial compounds profiled under consistent laboratory workflows, provides a substantial Gram-negative reference resource and reduces technical variability for cross-modality comparison.

Across all analyses, proteomics emerged as the most informative single modality. Whereas chemical structure fingerprints performed well for scaffolds already represented in the training set, they provided limited guidance for structurally distinctive compounds. Proteome-level responses, by contrast, enabled correct classification even when structural similarity was low, underscoring the value of protein-level data for capturing pathway-specific perturbations that are not evident from structure alone. However, the mechanistic information contained in proteomics data is distributed and not directly interpretable: classical enrichment analyses failed to recover the MoA for several classes in our dataset, indicating that machine learning is required to extract discriminative signatures from these high-dimensional profiles. This reliance on proteome-level signals rather than structural similarity is central for discovery efforts, where the priority is to predict modes of action for structurally novel compounds.

The text–feature matching formulation provided a practical solution to the limited sample size typical of mechanism-focused studies. Pairing biochemical fingerprints with augmented mechanistic descriptions expanded the training space without altering the underlying measurements, improving performance for both gradient-boosting and neural network models. As expected, the effect was stronger for the neural network, which is more dependent on larger training sets than boosting approaches[54]. Such augmentation strategies may therefore be broadly useful when experimental profiling is costly or biological replicates are limited.

Distinguishing correct predictions from model overconfidence remains a central challenge in MoA inference. The uncertainty estimator developed here addresses this by integrating replicate stability, ensemble variance, augmentation-derived disagreement, and probability-based metrics into a single score. Its behavior on structurally novel compounds illustrates the value of this approach: despite its comparatively high top-class prediction score, nitroxoline was flagged as uncertain because entropy- and variance-related signals remained elevated across perturbations, whereas cystobactamid remained below the uncertainty threshold despite a lower absolute prediction score because its entropy and replicate-variability signals were more consistent. These examples highlight the limits of probability-based confidence measures and show how stability-related features can help flag candidates with putatively novel modes of action rather than forcing assignment to known classes.

The external data experiments further illustrate both the strengths and the limits of proteomics-based MoA prediction. To avoid structural shortcuts, these analyses were intentionally conducted as a proteomics-only transfer variant of MAPPER. Raw protein abundance profiles generated on a different mass spectrometer or in another laboratory did not yield confident predictions, showing that absolute intensities are strongly shaped by platform- and workflow-specific effects. In contrast, significance-based filtering restored predictive performance across these heterogeneous datasets, indicating that differential protein responses form a more portable representation of antibiotic action than raw intensities.

The Subanovic dataset provided a particularly stringent test because antibiotics were applied at subinhibitory concentrations, yielding weak proteomic perturbations for most compounds. Consistent with this limited signal, significance-filtered inference produced a confident MoA prediction only for imipenem, the sole compound with a clear proteomic response, while the remaining antibiotics remained appropriately uncertain. Thus, when sufficient differential signal is present, significance-based proteomic features can support MoA prediction even outside the original training workflow without forcing assignments where evidence is weak.

Several limitations provide directions for future development. All measurements were conducted in *E. coli* under a single growth condition; extending the framework to additional Gram-negative species or environmental contexts may reveal species- or condition-specific signatures relevant for translation. Although the present work demonstrates clear gains over structure-only baselines, it does not provide a direct empirical comparison to other high-dimensional profiling modalities such as transcriptomics, metabolomics, or imaging, because matched datasets were not available for the compound panel studied here. Shared compound benchmarks would enable both fairer method comparison and testing of whether these complementary readouts can be integrated synergistically. Although significance filtering mitigated cross-laboratory differences in the datasets tested here, incorporating independent proteomics datasets during training would provide a stronger test of generalizability and dataset-specific technical effects. Expanded coverage of chemical diversity, particularly for underrepresented mechanistic classes, would further strengthen model generalization and novelty detection.

More broadly, this work serves as a proof-of-concept for proteomics-based predictive modeling in bacterial systems. While traditional antibiotic studies rely on lethal or growth-inhibitory phenotypes, proteome-level data capture non-fatal cellular responses that are inaccessible to classical screening readouts. As a result, similar approaches could be applied to pathways and phenotypes not tied to viability, such as virulence factor regulation, host-interaction mechanisms, persistence and stress-response pathways, as well as antibiotic resistance prediction. Predictive models trained on such datasets may help identify anti-virulence compounds or modulators of bacterial physiology that fall outside the scope of conventional antibiotic discovery yet remain relevant for therapeutic development. With the dataset provided in this study we aim to lay a foundation of proteomics data that can be extended and improved upon by future research endeavors. Because the underlying training data and feature representations are provided, MAPPER can be retrained on the full multimodal input space or adapted to reduced modality subsets matched to a user’s available data.

Together, this work defines MAPPER as a final multimodal MoA inference framework that combines a fixed predictor with uncertainty-aware decision support, while showing that a proteomics-only transfer variant retains mechanistic utility under external domain shift. The dataset and modeling approach presented here enable reproducible benchmarking, facilitate comparative development of computational tools and support more efficient exploration of chemical and mechanistic space in early discovery.

## Methods

### Bacterial strain and culturing

*E. coli* (strain K-12) was grown in LB medium at 37 ^°^C with shaking at 200 rpm. Unless noted otherwise, experiments were performed with solvent-only controls (typically DMSO).

### Phenotypic assays

#### Minimal inhibitory concentration (MIC)

Antibacterial activity against *E. coli* was determined using a broth microdilution MIC assay in 96-well plates (transparent Nunc flat-bottom, Thermo Fisher Scientific). The final bacterial inoculum was 5 × 10^5^ CFU mL^−1^. A 0.5 McFarland suspension of an overnight culture was diluted 1:100 in fresh LB medium, and 50 µL of this cell suspension was added to wells containing 50 µL of serially diluted compound solutions prepared at 2% solvent (1% final solvent concentration). DMSO was used as the standard solvent, while a small number of compounds required alternative stock solvents or conditions as listed in Supplementary Table 3; for compounds with very high MIC values, the final DMSO content exceeded 1% up to 2.5% to allow sufficient stock dilution. Each plate included untreated growth controls and an ampicillin reference row to ensure inter-plate consistency. After 24 h incubation at 37 ^°^C and 200 rpm, optical density at *λ* = 600 nm was measured using a Tecan Infinite^®^ 200 Pro M Nano reader. The MIC was defined as the lowest concentration showing no visible growth (OD_600_ ≤ 0.1). All compounds were assessed in technical triplicates.

#### Growth dynamics

Growth assays were performed at antibiotic concentrations corresponding to 1 ×, 5 ×, 10 ×, and, when required, with higher factors up to 50 × the MIC. An overnight culture was diluted in LB medium to an OD_600_ of 0.05, and 6 mL of this suspension were transferred into 14 mL round-bottom culture tubes. Cultures were incubated at 37 ^°^C and 200 rpm until reaching an OD_600_ of 0.4–0.6 (typically within 2 h), after which they were treated with the indicated antibiotic concentrations or solvent-only controls, in technical duplicates. OD_600_ was recorded hourly for 8 h to monitor growth dynamics using a BioPhotometer model 6131 (*Eppendorf*). After the 8 h measurement, 2 mL of fresh LB medium were added to each culture, and a final OD_600_ reading was collected after 24 h of incubation.

### Mass spectrometry

#### Full Proteome Analysis

The protocol was adapted from Köllen *et al*.[55]. A subset of the proteomics measurements generated within the present study—ampicillin, tetracycline, nitrofurantoin, nitrofurazone, and D8-treated samples—was also included in Köllen *et al*.[55] and published there; these compounds are marked in Supplementary Table 3. Overnight cultures of *E. coli* K-12 were diluted to an initial OD_600_ of 0.05 in 5.5 mL LB medium and incubated for 2 h at 37 ^°^C with shaking at 200 rpm. Optical density was determined after dilution (1:1). When cultures reached OD_600_ ≈ 0.5, cells were treated either with antibiotics or the corresponding solvent controls; the compound-specific treatment concentrations, MIC factors, and matched controls used for proteomics sample generation are listed in Supplementary Table 3. Cultures were further incubated for 1 h at 37 ^°^C with shaking at 200 rpm. Following incubation, cultures were normalized by dilution with PBS to match the lowest OD_600_.

1 mL of each normalized culture was harvested by centrifugation (6000 g, 5 min, 4 ^°^C) and washed once with ice-cold PBS. Cell pellets were resuspended in 150 µL PBS supplemented with 0.5 % SDS and 1 % Triton X-100. Cell disruption was achieved using a 10 s sonication pulse at 30 % intensity (Sonopuls HD 2070, *Bandelin*), followed by bead beating in bead-mill tubes containing 0.1 mm zirconia beads. Bead beating was performed for three cycles of 30 s at 6500 rpm with 30 s cooling intervals between cycles using a Precellys 24 homogenizer (*Bertin Technologies*).

Cell lysates were centrifuged at 10 000 g for 10 min, after which the supernatant was transferred to fresh microcentrifuge tubes and centrifuged again at 21 000 g for 30 min at room temperature. The resulting supernatants were transferred into LoBind tubes (*Eppendorf*), and protein concentrations were determined using a BCA assay (Roti Quant, *Carl Roth*).

For downstream processing, equal protein amounts (50 µg in 80 µL) were transferred into a V-bottom 96-well polypropylene plate (Greiner, cat. 651201). Reduction and alkylation were carried out by adding 3 µL of a 1:2 mixture of tris(2-carboxyethyl)phosphine (TCEP) and iodoacetamide (IAA) (each 500 mM), followed by incubation for 15 min at 950 rpm at room temperature. Residual IAA was quenched by adding 2 µL of 500 mM dithiothreitol (DTT). Subsequently, a 1:1 mixture of hydrophobic and hydrophilic carboxylate-coated magnetic beads (*Cytiva*, cat. 65152105050250 and 45152105050250), previously washed three times with water, was added (10 µL per well). Protein precipitation was induced by adding 150 µL ethanol.

All subsequent liquid handling steps were performed using a Hamilton Microlab Prep robotic workstation. Samples were shaken for 5 min at 500 rpm at room temperature, after which magnetic beads were collected using a 96-ring magnet (Magnum FLX, *Alpaqua*). Supernatants were removed slowly (20 µL s^−1^) to minimize bead loss. The beads were washed three times with 180 µL 80 % ethanol and once with 180 µL acetonitrile, with off-magnet shaking between washing steps (1 min, 800 rpm, room temperature).

Protein digestion was carried out overnight at 37 ^°^C in 100 µL 50 mM TEAB containing 1 µL sequencing-grade trypsin (trypsin:protein ratio 1:100, 0.5 µg µL^−1^; *Promega*). Digestion was performed with shaking at 800 rpm under a heated lid with the plate tightly sealed. Peptides were eluted by adding 50 µL 3 % formic acid and subsequently desalted using two-disk styrenedivinylbenzene reverse-phase sulfonate (SDB-RPS) StageTips (Empore, *3M*). StageTips were equilibrated with 150 µL wash buffer 1 (1 % TFA in isopropanol)[56]. Samples were loaded by centrifugation (10 min, 500 g), followed by washing with buffer 1 (30 min, 800 g) and buffer 2 (0.2 % TFA in water; 30 min, 800 g). Peptides were eluted using 50 µL elution buffer (1 % NH_3_, 80 % acetonitrile) by centrifugation (5 min at 300 g followed by 800 g). Eluted peptides were dried in a centrifugal evaporator (Concentrator Plus, *Eppendorf*) and subsequently reconstituted in 1 % formic acid. For LC-MS/MS analysis, 3 µL of each sample was injected into a *Bruker* timsTOF Pro instrument operated in DIA mode. All experimental conditions were analyzed in four independent biological replicates.

#### LC-MS/MS Measurements timsTOF Pro

The protocol was adapted from Köllen *et al*.[55]. Peptide separation and mass spectrometric analysis were performed on an UltiMate 3000 nano-HPLC system (*Dionex*) coupled to a timsTOF Pro mass spectrometer (*Bruker*) equipped with a CaptiveSpray nano-electrospray ion source and a *Sonation* column oven. Samples were initially loaded onto a trap column (Acclaim PepMap 100 C18, 75 µm ID × 2 cm, 3 µm particle size, *Thermo Fisher Scientific*) and washed for 7 min using solvent A (0.1 % formic acid in water) at a flow rate of 5 µL min^−1^.

Peptides were subsequently transferred to an analytical column (Aurora C18, 25 cm × 75 µm, 1.7 µm particle size, *IonOpticks*) and separated using a gradient of solvent B (0.1 % formic acid in acetonitrile) at a constant flow rate of 400 nL min^−1^. The gradient program consisted of 5–28 % solvent B over 28 min, followed by an increase to 40 % B over 6 min. A high-organic wash at 95 % B was applied for 6 min, after which the column was re-equilibrated at 5 % B for 10 min.

Mass spectrometric acquisition was performed in dia-PASEF mode. Ion mobility separation was achieved using a dual TIMS analyzer with accumulation and ramp times of 100 ms. MS1 scans covered an ion mobility range of 0.60 V s cm^−2^ to 1.60 V s cm^−2^. Fragmentation was performed over an m/z range of 400–1201 using an ion mobility window spanning 0.60 V s cm^−2^ to 1.43 V s cm^−2^.

Each dia-PASEF scan comprised two ion mobility isolation windows with a width of 26 m/z. The entire mass range was covered using 32 windows with 1 m/z overlaps, corresponding to 16 dia-PASEF scans per MS1 cycle and resulting in an overall cycle time of approximately 1.80 s. The full DIA-PASEF window scheme, including ion mobility ranges and scan boundaries, is listed in Supplementary Table 4. Collision energy was adjusted linearly from 59 eV at 1/K_0_ = 1.3 V s cm^−2^ to 20 eV at 1/K_0_ = 0.85 V s cm^−2^. Calibration of the TIMS elution voltage was performed using reference ions (m/z 622, 922, and 1222) from the Agilent ESI-L Tuning Mix introduced into the CaptiveSpray inlet filter to obtain accurate reduced ion mobility coefficients (1/K_0_).

#### LC-MS/MS Measurements Orbitrap Eclipse

For the experiment using a different mass spectrometer, the protocol was adapted from Bottlinger *et al*.[57]. Peptide analysis was performed using a Vanquish Neo UHPLC system (*Thermo Fisher Scientific*) coupled to an Orbitrap Eclipse Tribrid mass spectrometer (*Thermo Fisher Scientific*). The UHPLC system was equipped with a PepMap Neo C18 trap cartridge (5 µm, 300 µm × 5 mm, *Thermo Fisher Scientific*) and operated in trap-and-elute injection mode, in which samples were first loaded onto the trap column prior to chromatographic separation.

Chromatographic separation was performed on an Aurora Ultimate separation column (3rd generation, 25 cm, *IonOpticks*) maintained at 40 ^°^C and connected to a Nanospray Flex ion source (*Thermo Fisher Scientific*). The system was operated at a flow rate of 400 nL min^−1^ using solvent A (0.1 % formic acid in water) and solvent B (0.1 % formic acid in acetonitrile). Peptides were separated using a 45 min gradient starting from 5 % to 28 % solvent B over 30 min, followed by an increase to 35 % B within 5 min. A high-organic wash step at 90 % solvent B was then applied for 10 min. Column washing and equilibration were performed using the instrument’s fast equilibration mode (equilibration factor 3) at 5 % solvent B. Trap column cleaning and equilibration were carried out using fast wash and equilibration in combination with zebra wash (two wash cycles with automatic equilibration factor).

Mass spectrometric acquisition was performed on the Orbitrap Eclipse using internal real-time mass calibration with a user-defined lock mass at m/z 445.12003. Data was acquired in data-independent acquisition (DIA) mode. Full MS scans were recorded in the Orbitrap at a resolution of 60 000 with an AGC target of 4 × 10^5^ and a maximum injection time of 100 ms across an m/z range of 400–1000. Fragment ion spectra (MS2) were acquired at a resolution of 15 000 with an AGC target of 1 × 10^6^ and a maximum injection time of 40 ms. Precursor isolation was performed in the quadrupole using windows of 10 m/z with 1 m/z overlap covering an m/z range of 145–1450. Fragmentation was achieved by higher-energy collision-induced dissociation (HCD) using a normalized collision energy of 30 %. Instrument control and data acquisition were performed using Thermo Scientific Foundation software (version 3.1sp9) together with Xcalibur (version 4.6). Raw files generated by the Orbitrap Eclipse were converted to mzML format using the MSConvert tool (version 3.0.21193-ccb3e0136) from the ProteoWizard software suite (version 3.0.21193, 64-bit).

#### Proteomics quantification and differential analysis

Raw mass spectrometry data (or the transformed mzML files for Eclipse data) was processed in DIA-NN (library-free mode, version 1.8.1)[58]. For library generation, the UniProt reference proteome for *E. coli* K-12 (proteome ID: UP000000625; taxon ID: 83333; downloaded on 2024/07/16) was used. Protein-level quantification tables were further processed in Perseus (version 2.1.3.0): LFQ intensities were log_2_-transformed; protein groups were retained if at least three values were present in at least one treatment group. For significance-gated proteomics data, antibiotic-treated conditions were compared against matched solvent controls using two-sample Student’s t-tests with permutation-based multiple-testing correction (FDR = 0.05).

#### Proteomics representations used for modeling

For modeling on the in-house dataset, proteomics input matrices were constructed from the processed Perseus exports by retaining quantitative intensity columns together with gene annotations. Compound were profiled in four biological replicates, except doripenem, azithromycin, cerulenin, and squalamine, which each had 3 biological replicates due to workflow issues. Gene identifiers were expanded when multiple gene names were assigned to a protein group, and a unified gene-by-sample matrix was assembled over the *E. coli* K-12 gene universe (UniProt FASTA). Missing values were set to zero and all entries were coerced to numeric values, yielding a 4,401-feature proteomics vector per sample (`X_proteomics.pkl`).

For visualization of the proteomics landscape in Figure 2d, a two-dimensional unsupervised UMAP embedding was computed from the sample-by-gene proteomics matrix using Uniform Manifold Approximation and Projection (UMAP)[59].

For cross-platform and cross-laboratory analyses, we additionally used a significance-filtered proteomics representation in which replicate intensities for proteins not meeting a per-protein significance criterion relative to controls were set to zero (`X_proteomics_sig.pkl`). Unless stated otherwise, significance filtering used a threshold of 1.3 on the − log_10_(*p*) scale (approximately *p <* 0.05).

### Auxiliary modalities

#### Chemical structure features

Chemical structure features were computed from SMILES strings as Morgan fingerprints (radius 2; 2,048 bits) using RDKit[60].

For visualization of structural relationships in Figure 2c, a two-dimensional unsupervised UMAP embedding was computed from the compound-level ECFP vectors using Uniform Manifold Approximation and Projection (UMAP)[59].

#### Cell-painting proxy embeddings (InfoAlign)

To provide an auxiliary morphological–transcriptional proxy representation, we computed 300-dimensional embeddings from SMILES using a pretrained InfoAlign model [34].

#### MIC and growth curve feature vectors

MIC profiles were mapped onto a two-fold dilution series (800 to 0.001), missing concentrations were filled by nearest measured values, and features were min–max scaled per compound to generate a 20-dimensional MIC vector. Growth curve features were derived by selecting the lowest antibiotic concentration condition whose OD at the final time point (24 h) was below 50% of the solvent control (or the highest tested concentration if none met this criterion). The selected curve was multiplied by the MIC factor used for that condition and summarized as a 10 time-point feature vector.

### MoA text descriptions and embeddings

For each MoA class, a mechanistic description (approximately 200 words) was curated and expanded into ten variants per augmentation regime. Sentence-level and word-level augmentations were generated by random shuffling (with fixed seeds); semantic augmentation consisted of independently rewritten paraphrases intended to preserve mechanistic content while varying phrasing. These semantic paraphrases were generated with ChatGPT (OpenAI) [61]. All generated outputs were manually reviewed, screened for factual consistency and redundancy, and edited where necessary before downstream analysis. MoA descriptions were embedded using transformer encoders (PubChemDeBERTa-augmented, BERT-base-uncased, BioBERT v1.1); exact Hugging Face model revisions (commit SHAs) and embedding settings are reported in the Supplementary Information [39–41]. The augmentation/encoder benchmark (Figure 3) evaluated all combinations, whereas downstream analyses (Figures 4–6) used a fixed semantic+BioBERT configuration.

### Learning task formulation

MoA prediction was formulated as a compound–MoA pairwise binary classification task. For each experimental sample, we constructed rows by crossing the compound-level feature vector (selected modalities) with each candidate MoA text embedding. The binary target was defined as 1 when the candidate MoA matched the true MoA label and 0 otherwise. For multiclass readouts, pairwise scores were aggregated across text variants and the highest-scoring MoA was selected. No-text baselines operated directly on compound-level features without pairwise expansion.

### Models

The primary deployment model was TabM[42]. Comparative modeling experiments additionally evaluated a PyTorch multilayer perceptron[62], a scikit-learn multilayer perceptron[63], XGBoost[64], LightGBM[65] and a scikit-learn random-forest classifier. TabM was trained for up to 100 epochs (batch size 256) with early stopping (patience 3), learning rate 1 × 10^−4^ and weight decay 3 × 10^−4^. Detailed usage of the other models is shown in the supplementary information.

### Evaluation protocol and preprocessing

To avoid leakage between chemically identical compounds, all cross-validation splits were generated at the compound level. Figure-linked benchmarking used stratified five-fold cross-validation with five independent fold partitions (base seed = 42), and all pairwise-expanded rows inherited the split assignment of their parent compound.

Within each training fold, features were standardized using `StandardScaler` fit on the training data and applied to held-out data. Random seeds were set for Python, NumPy and PyTorch. For uncertainty experiments, we performed LOO (held-out compound) and LOCO (held-out MoA class) evaluations with five seed repetitions.

### Performance metrics and statistical analyses

Binary performance was summarized using PR AUC, ROC AUC and Matthews correlation coefficient. Multiclass comparisons used macro-averaged F1 and one-vs-rest macro PR AUC and macro ROC AUC.

For model and augmentation comparisons across repeated cross-validation cycles, we followed guidelines presented by Ash et al [66], using paired, repeated-measures statistical tests as appropriate for each analysis (including repeated-measures ANOVA with Tukey-style post hoc comparisons, and Friedman tests with post hoc paired comparisons and multiple-testing correction). A detailed per-figure accounting of statistical tests, multiple-testing correction, and effect-size estimation is provided in the Supplementary Information.

### Uncertainty estimation

Although training is pairwise binary (match vs non-match for each sample–candidate-MoA row), uncertainty is computed from an aggregated multiclass readout. For compound *i* and candidate class *c*, we aggregate binary match scores over all available perturbation-level predictions, including biological replicates, text augmentations, and seed repetitions:

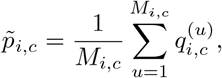

where 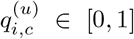 is the binary match probability for perturbation index *u* and candidate class *c*, and *M*_*i,c*_ is the number of available perturbation-level predictions contributing to that compound–class pair. The resulting class-score vector 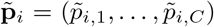 is used for top-class selection and margin-based uncertainty features. For entropy and disagreement features, we write 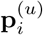 for the normalized class distribution of perturbation *u* and 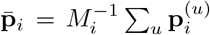 for its perturbation average.

We define the top score and margin as

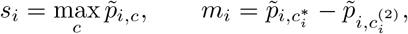

where 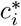 and 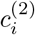 denote the highest- and second-highest-probability classes, respectively. The uncertainty feature set combines standard confidence- and information-theoretic quantities, including maximum class score, predictive entropy, expected entropy, mutual-information-based disagreement and ensemble-style variation measures[67–70], together with replicate- and perturbation-stability features introduced here for the current experimental setting. Concretely, PredScore = *s*_*i*_ and Margin = *m*_*i*_; PredEntropy is the entropy of the perturbation-averaged class distribution, ExpEntropy is the perturbation-averaged entropy, and EpistemicMI = PredEntropy − ExpEntropy. MeanPairKL denotes the mean sampled pairwise Kullback–Leibler divergence between perturbation-level class distributions; KConsistency is the modal class agreement fraction across perturbations and VariationRatio = 1 − KConsistency. VarMax is the variance of the perturbation-level maximum class probability, AugVar the augmentation-induced class-score variance for the predicted class, SeedStd the seed-level variability of the predicted-class score, and BioVar the variance of predicted-class scores across biological replicates. Writing 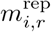 for the top-vs-second margin computed within replicate *r*, RepMarginStd = 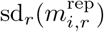 and RepMarginIQR 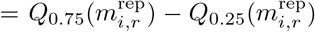, while RepAgree is the fraction of replicates whose top class matches the compound-level top class. Detailed mathematical definitions of all derived uncertainty features are provided in the Supplementary Information.

We benchmarked six uncertainty estimators (Fig. 5b-c), all converted to “higher = more uncertain” scores:

1. **Low max score**: 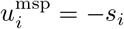.
2. **High predictive entropy**: 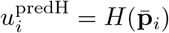, where *H*(**p**) = − ∑_*c*_ *p*_*c*_ log *p*_*c*_ and 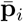 is the perturbation-averaged class distribution.
3. **High expected entropy**: 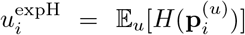, i.e., entropy averaged over perturbations *u*.
4. **Z-Gate (global)**: a rule-based score combining (i) positive standardized deviations of selected risk signals and (ii) a low-margin penalty:

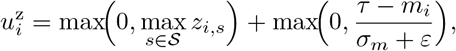

with *z*_*i,s*_ = *d*_*s*_(*x*_*i,s*_ − *µ*_*s*_)*/*(*σ*_*s*_ + *ε*), sign *d*_*s*_ ∈ {−1, +1} set by risk direction, and 𝒮 = {MeanPairKL, PredEntropy, EpistemicMI, BioVar, RepMarginStd}. In the global variant, (*τ, µ*_*s*_, *σ*_*s*_) are estimated once from correct LOO development samples.
5. **Z-Gate (class-conditional)**: same functional form as above, but *τ, µ*_*s*_, and *σ*_*s*_ are estimated per predicted class and shrinkage-regularized toward global estimates.
6. **Meta-model (LR+Poly)**: calibrated logistic regression on a multifeature risk vector including score/margin, entropy/disagreement, augmentation/seed variance, and replicate-consistency features. Before fitting, PredScore and Margin were sign-inverted so that larger values uniformly indicate greater prediction risk. The pipeline is

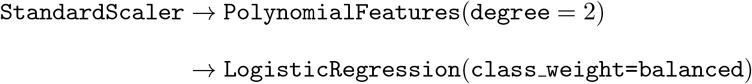

wrapped in `CalibratedClassifierCV` (sigmoid, cv=3), and outputs 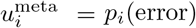.

All six methods were threshold-calibrated to the same in-distribution target false-positive rate (ID-FPR = 0.10) using only correct LOO development predictions. For a method score *u*, the uncertainty threshold is the (1 − ID-FPR)-quantile of *u* on LOO-correct samples; a compound is flagged uncertain if *u*_*i*_ ≥ *t*. This harmonized calibration enables a direct comparison of error-recall performance at matched in-distribution conservativeness.

For transfer settings (Figure 6), we additionally applied conservative floor-based overrides before the final uncertainty call. The score floor for the top class probability is

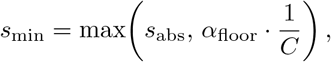

where *s*_abs_ is the absolute minimum score floor (`abs_min_score_floor`), *α*_floor_ is the relative score-floor factor (`score_floor_factor`), and *C* is the number of candidate classes. A second floor is applied to the decision margin,

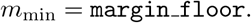

If either *s*_*i*_ *< s*_min_ or *m*_*i*_ *< m*_min_, the sample is force-flagged as uncertain (equivalently, *p*_error_ = 1). Intuitively, these floors prevent confident calls when both top-class support and class separation are too weak.

### External evaluation and domain-shift handling

To assess robustness under domain shift, we applied models to proteomics measured on a different instrument and to an external dataset generated in an independent laboratory. For these evaluations, we used the significance-filtered proteomics representation and applied uncertainty estimation with additional floor-based rules (absolute minimum score floor 0.05; score floor factor 0.1; margin floor 0.01) to conservatively flag low-signal cases as uncertain. For Figure 6 transfer, the uncertainty model weights were kept fixed and only the uncertainty threshold was recalibrated on development LOO/LOCO metadata generated under the semantic+proteomics setting; no external or eclipse test labels were used for this threshold adaptation.

### Usage of generative AI

Generative AI tools (ChatGPT and Codex) were used to assist with code drafting, code editing and manuscript wording; all resulting code, analyses and text were reviewed, validated and revised by the authors, who take full responsibility for the final content.

## Data availability

The mass spectrometry proteomics dataset generated in this study has been deposited to the ProteomeXchange Consortium via the PRIDE [71] partner repository under accession PXD076144. All processed feature matrices and derived embeddings used in this study are available at https://github.com/sieber-lab/MAPPER.

## Code availability

All code used for data processing, model training, evaluation, and figure generation is available at https://github.com/sieber-lab/MAPPER. The InfoAlign implementation used in this study is available at https://github.com/liugangcode/InfoAlign[34].

## Acknowledgements

The authors thank Merck KGaA Darmstadt for their generous support with the Merck Future Insight Prize 2020. This project was also cofunded by the European Union (ERC, breakingBAC, 101096911). Views and opinions expressed are however those of the author(s) only and do not necessarily reflect those of the European Union or the European Research Council. Neither the European Union nor the granting authority can be held responsible for them. The authors thank M. Köllen for providing the compounds D8, D12, and Debio-1452-NH2. Joshua Hesse thanks M. Schuh and A. Daniluk for valuable discussions and feedback on this work.

## Author contributions

Jo.H. and S.A.S. conceived the project. Jo.H. developed the methodologies and the codebase, performed data analysis, and wrote the manuscript. Jo.H. and L.L. performed microbiology experiments. D.S. and L.L. performed proteomics experiments. L.R.G. performed oganic synthesis. Je.H. and R.M. contributed natural products and proofread the manuscript.

## Additional information

Supplementary information accompanies this paper. Correspondence and requests for materials should be addressed to S.A.S..

## Extended Data

**Extended Figure 1.**
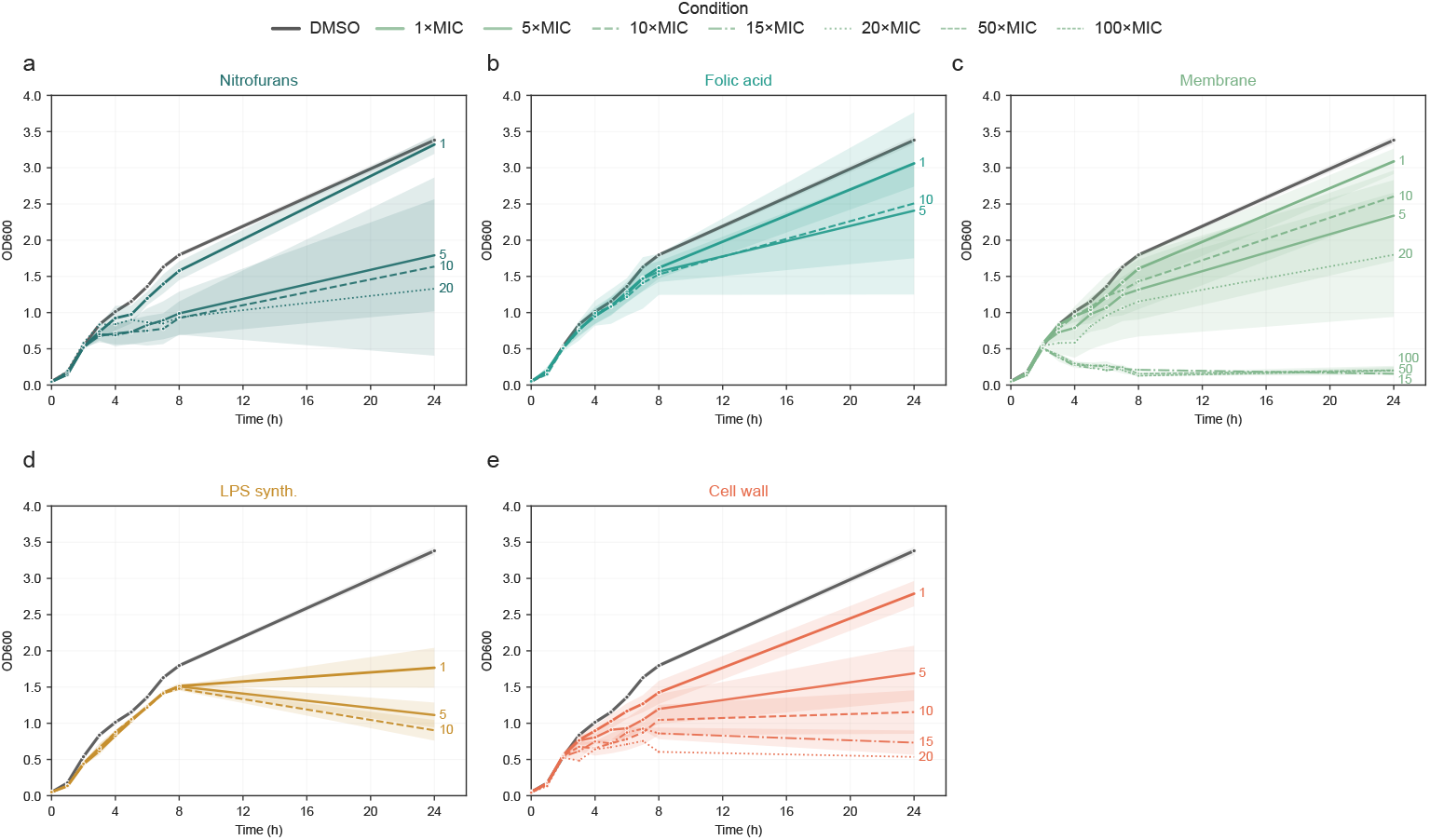
Growth curve results per MoA class: Growth curve effects for all remaining MoA classes. Per class, growth curve values for each timepoint (one measurement per hour for 0-8 h and final measurement at 24 h) were averaged across all class members. The shaded areas show the SEM band for each concentration curve. MIC factor is represented by the type of line used, and is also shown at the next to the 24 h point of the respective curve.

**Extended Figure 2.**
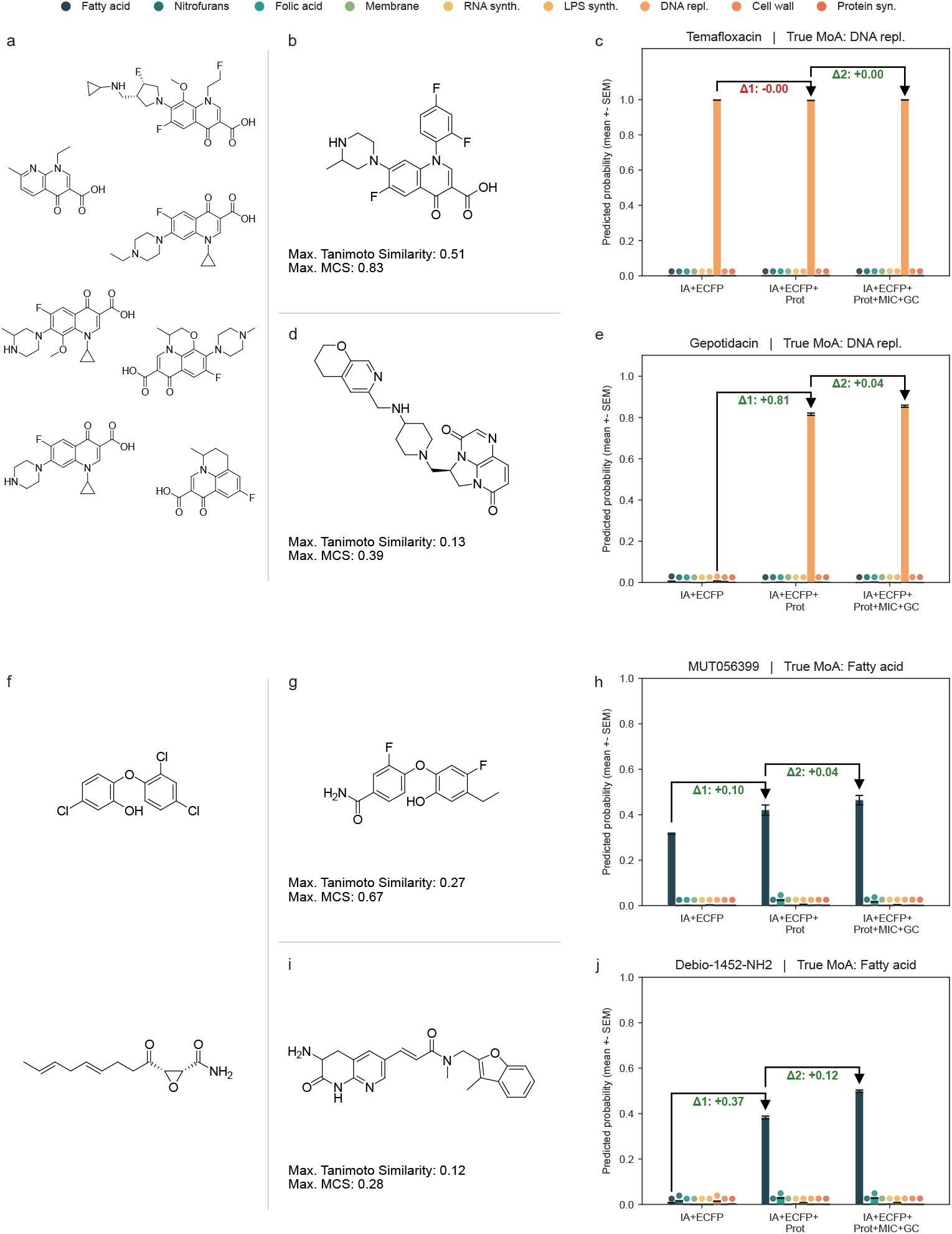
Structure-dependent Feature-effects: (a) Chemical structures of the remaining DNA synthesis inhibitors. (b, d) Structure of temafloxacin & gepotidacin, maximum Tanimoto similarity and maximum common substructure score compared to remaining class members. (c, e) Temafloxacin and gepotidacin predicted MoA probabilities using InfoAlign and chemical structure fingerprints vs adding proteomics vs all feature modalities. Bars show mean ± SEM across *n* = 200 replicate predictions per MoA; colored dots denote MoA categories. Arrows and labels indicate the delta in mean predicted probability for the true MoA due to addition of proteomics (Δ1) and MIC & GC data (Δ2). (f) Chemical structures of the remaining fatty acid synthesis inhibitors. (g, i) Structure of MUT056399 & Debio-1452-NH_2_, maximum Tanimoto similarity and maximum common substructure score compared to remaining class members. (h, j) MUT056399 and Debio-1452-NH_2_ predicted MoA probabilities using InfoAlign and chemical structure fingerprints vs adding proteomics vs all feature modalities. Bars show mean ± SEM across *n* = 200 replicate predictions per MoA; colored dots denote MoA categories. Arrows and labels indicate the delta in mean predicted probability for the true MoA due to addition of proteomics (Δ1) and MIC & GC data (Δ2).

**Extended Figure 3.**
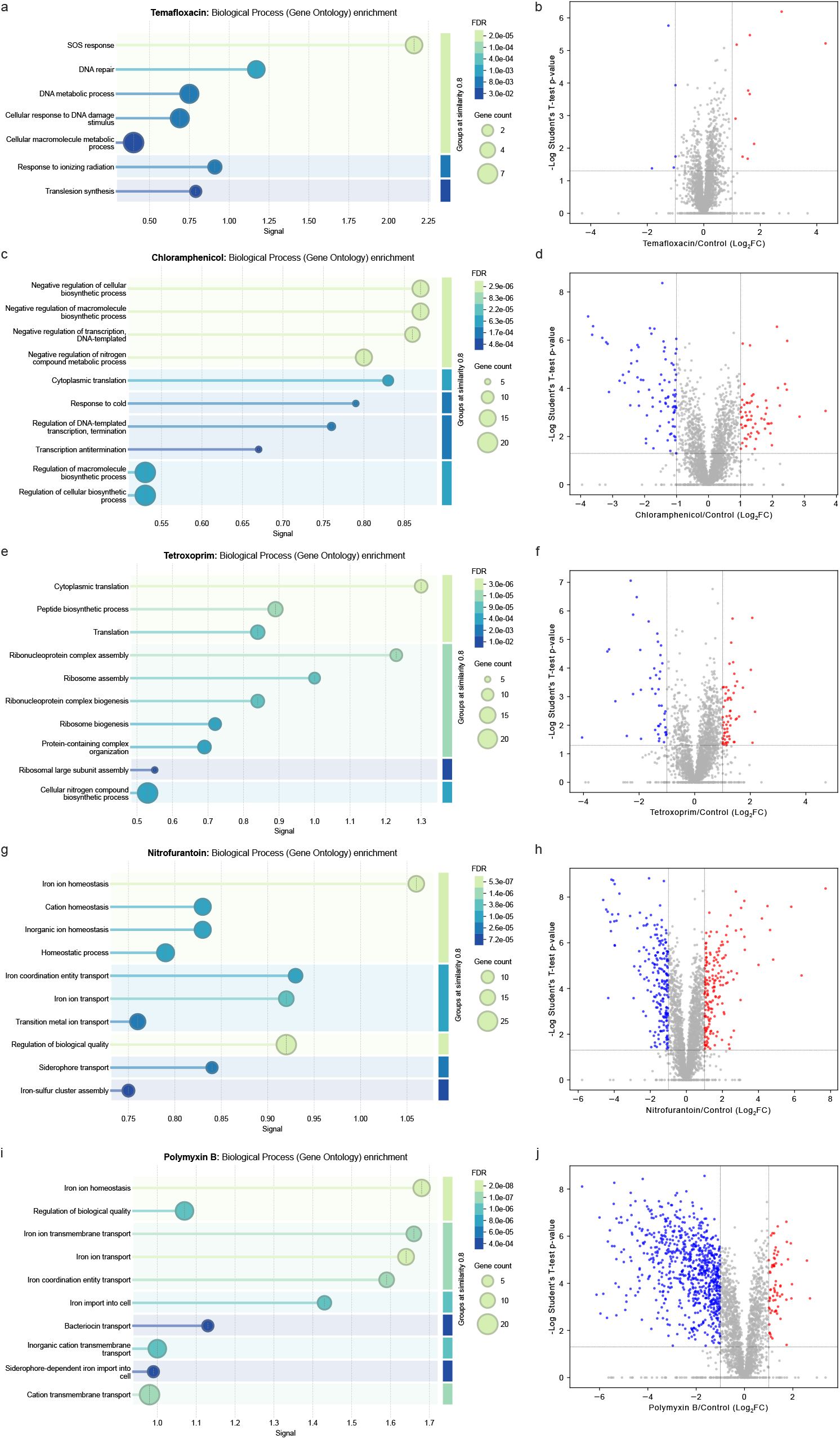
Proteome-based pathway enrichment analysis: Gene Ontology Biological Process enrichments generated via the STRING web interface [43] (left) and corresponding volcano plots of differential protein abundances (right) for representative inhibitors of DNA synthesis (a–b), protein synthesis (c–d), folate metabolism (e–f), nitrofuran antibiotics (g–h) and membrane integrity (i–j). In the volcano plots, vertical black lines mark ±1-fold change and the horizontal line marks the statistical significance threshold (p = 0.05).

**Extended Figure 4.**
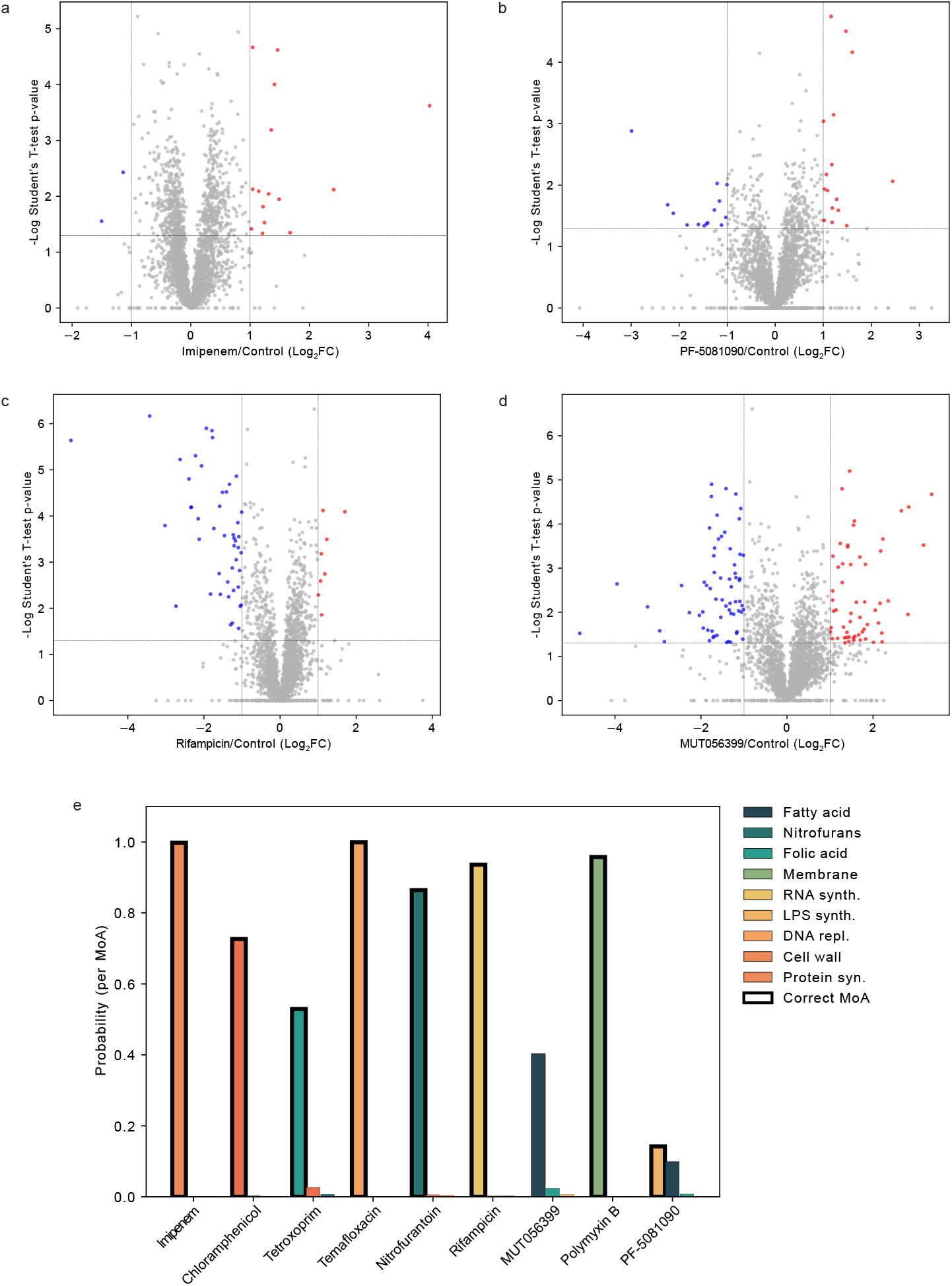
ML–based MoA classification from proteomic responses despite absent pathway-level enrichment: Volcano plots of differential protein abundances for representative inhibitors of cell wall synthesis (a), LPS synthesis (b), RNA synthesis (c), and fatty acid synthesis (d). Gene Ontology Biological Process enrichment identified 0 enriched processes via the STRING web interface [43] for these samples. In the volcano plots, vertical black lines mark ±1-fold change and the horizontal line marks the statistical significance threshold (p = 0.05). (e) Top three MoA prediction probabilities for each of the nine MoA representatives using proteomics as the only feature set.

## References

[1] Kwon, J. H. & Powderly, W. G. The post-antibiotic era is here. Science 373, 471–471 (2021). URL https://www.science.org/doi/10.1126/science.abl5997.

[2] Reardon, S. WHO warns against ‘post-antibiotic’ era. Nature nature.2014.15135 (2014). URL https://www.nature.com/articles/nature.2014.15135.

[3] Miethke, M. et al. Towards the sustainable discovery and development of new antibiotics. Nature Reviews Chemistry 5, 726–749 (2021). URL https://www.nature.com/articles/s41570-021-00313-1.

[4] Payne, D. J., Gwynn, M. N., Holmes, D. J. & Pompliano, D. L. Drugs for bad bugs: confronting the challenges of antibacterial discovery. Nature Reviews Drug Discovery 6, 29–40 (2007). URL https://www.nature.com/articles/nrd2201.

[5] Gargate, N., Laws, M. & Rahman, K. M. Current economic and regulatory challenges in developing antibiotics for Gram-negative bacteria. npj Antimicrobials and Resistance 3, 50 (2025). URL https://pmc.ncbi.nlm.nih.gov/articles/PMC12159177/.

[6] Beyer, P. & Paulin, S. The Antibacterial Research and Development Pipeline Needs Urgent Solutions. ACS Infectious Diseases 6, 1289–1291 (2020). URL 10.1021/acsinfecdis.0c00044.

[7] Melchiorri, D., Rocke, T., Alm, R. A., Cameron, A. M. & Gigante, V. Addressing urgent priorities in antibiotic development: insights from WHO 2023 antibacterial clinical pipeline analyses. The Lancet. Microbe 6, None (2025). URL https://pmc.ncbi.nlm.nih.gov/articles/PMC11876093/.

[8] Breijyeh, Z., Jubeh, B. & Karaman, R. Resistance of Gram-Negative Bacteria to Current Antibacterial Agents and Approaches to Resolve It. Molecules 25, 1340 (2020). URL https://www.mdpi.com/1420-3049/25/6/1340. Publisher: Multidisciplinary Digital Publishing Institute.

[9] World Health Organization. 2021 antibacterial agents in clinical and preclinical development: an overview and analysis (World Health Organization, 2022).

[10] Romano, K. P. et al. Perturbation-specific transcriptional mapping for unbiased target elucidation of antibiotics. Proceedings of the National Academy of Sciences 121, e2409747121 (2024). URL https://www.pnas.org/doi/10.1073/pnas. 2409747121.

[11] Cunha, B. R. d., Zoio, P., Fonseca, L. P. & Calado, C. R. C. Technologies for High-Throughput Identification of Antibiotic Mechanism of Action. Antibiotics 10 (2021). URL https://www.mdpi.com/2079-6382/10/5/565.

[12] Mongia, M., Guler, M. & Mohimani, H. An interpretable machine learning approach to identify mechanism of action of antibiotics. Scientific Reports 12, 10342 (2022). URL https://www.nature.com/articles/s41598-022-14229-3. Publisher: Nature Publishing Group.

[13] Ioerger, T. R. et al. Identification of New Drug Targets and Resistance Mechanisms in Mycobacterium tuberculosis. PLOS ONE 8, e75245 (2013). URL https://journals.plos.org/plosone/article?id=10.1371/journal.pone.0075245. Publisher: Public Library of Science.

[14] A. Farha, M. & D. Brown, E. Strategies for target identification of antimicrobial natural products. Natural Product Reports 33, 668–680 (2016). URL https://pubs.rsc.org/en/content/articlelanding/2016/np/c5np00127g. Publisher: Royal Society of Chemistry.

[15] Porta, E. O. & Steel, P. G. Activity-based protein profiling: A graphical review. Current Research in Pharmacology and Drug Discovery 5, 100164 (2023). URL https://linkinghub.elsevier.com/retrieve/pii/S2590257123000123.

[16] Jin, Y., Jana, S., Abbasov, M. E. & Lin, H. Antibiotic target discovery by integrated phenotypic and activity-based profiling of electrophilic fragments. Cell Chemical Biology 32, 434–448.e9 (2025). URL https://www.sciencedirect.com/science/article/pii/S2451945625000339.

[17] Bond, A. N. et al. Reference-based chemical-genetic interaction profiling to elucidate small molecule mechanism of action in Mycobacterium tuberculosis. Nature Communications 16, 9673 (2025). URL https://www.nature.com/articles/s41467-025-64662-x. Publisher: Nature Publishing Group.

[18] Liu, C. et al. Deep learning-driven prediction of drug mechanism of action from large-scale chemical-genetic interaction profiles. Journal of Cheminformatics 14, 12 (2022). URL 10.1186/s13321-022-00596-6.

[19] Monfort-Lanzas, P., Rungger, K., Madersbacher, L. & Hackl, H. Machine learning to dissect perturbations in complex cellular systems. Computational and Structural Biotechnology Journal 27, 832–842 (2025). URL https://linkinghub.elsevier.com/retrieve/pii/S2001037025000583.

[20] Vora, N., Shah, S., Patel, P. & Shah, M. Artificial intelligence and multiomics in drug discovery: A deep learning-powered revolution. Cure & Care 1, 100011 (2025). URL https://www.sciencedirect.com/science/article/pii/S3051062725000112.

[21] Espinoza, J. L. et al. Predicting antimicrobial mechanism-of-action from transcriptomes: A generalizable explainable artificial intelligence approach. PLOS Computational Biology 17, e1008857 (2021). URL https://journals.plos.org/ploscompbiol/article?id=10.1371/journal.pcbi.1008857. Publisher: Public Library of Science.

[22] Senges, C. H. R. et al. Comparison of Proteomic Responses as Global Approach to Antibiotic Mechanism of Action Elucidation. Antimicrobial Agents and Chemotherapy 65, 10.1128/aac.01373–20 (2020). URL https://journals.asm.org/doi/10.1128/aac.01373-20. Publisher: American Society for Microbiology.

[23] Yu, Y. et al. Predictive Signatures of 19 Antibiotic-Induced Escherichia coli Proteomes. ACS Infectious Diseases 6, 2120–2129 (2020). URL https://pubs.acs.org/doi/10.1021/acsinfecdis.0c00196.

[24] Vincent, I. M., Ehmann, D. E., Mills, S. D., Perros, M. & Barrett, M. P. Untargeted Metabolomics To Ascertain Antibiotic Modes of Action. Antimicrobial Agents and Chemotherapy 60, 2281–2291 (2016). URL https://journals.asm.org/doi/10.1128/aac.02109-15. Publisher: American Society for Microbiology.

[25] Zampieri, M. et al. High-throughput metabolomic analysis predicts mode of action of uncharacterized antimicrobial compounds. Science Translational Medicine 10, eaal3973 (2018). URL https://www.science.org/doi/10.1126/scitranslmed.aal3973. Publisher: American Association for the Advancement of Science.

[26] Belenky, P. et al. Bactericidal Antibiotics Induce Toxic Metabolic Perturbations that Lead to Cellular Damage. Cell Reports 13, 968–980 (2015).

[27] Krentzel, D. et al. Deep learning recognises antibiotic modes of action from brightfield images (2025). URL https://www.biorxiv.org/content/10.1101/2025.03.30.645928v3. Pages: 2025.03.30.645928 Section: New Results.

[28] Liu, Y., Beyer, A. & Aebersold, R. On the Dependency of Cellular Protein Levels on mRNA Abundance. Cell 165, 535–550 (2016). URL https://www.cell.com/cell/abstract/S0092-8674(16)30270-7.

[29] Vogel, C. & Marcotte, E. M. Insights into the regulation of protein abundance from proteomic and transcriptomic analyses. Nature Reviews Genetics 13, 227– 232 (2012). URL https://www.nature.com/articles/nrg3185.

[30] Nonejuie, P., Burkart, M., Pogliano, K. & Pogliano, J. Bacterial cytological profiling rapidly identifies the cellular pathways targeted by antibacterial molecules. Proceedings of the National Academy of Sciences 110, 16169–16174 (2013). URL https://www.pnas.org/doi/10.1073/pnas.1311066110.

[31] Zoffmann, S. et al. Machine learning-powered antibiotics phenotypic drug discovery. Scientific Reports 9, 5013 (2019). URL https://www.nature.com/articles/s41598-019-39387-9.

[32] Bray, M.-A. et al. Cell Painting, a high-content image-based assay for morphological profiling using multiplexed fluorescent dyes. Nature Protocols 11, 1757–1774 (2016). URL https://www.nature.com/articles/nprot.2016.105.

[33] Tsakou, F., Jersie-Christensen, R., Jenssen, H. & Mojsoska, B. The Role of Proteomics in Bacterial Response to Antibiotics. Pharmaceuticals 13 (2020). URL https://www.mdpi.com/1424-8247/13/9/214.

[34] Liu, G. et al. Learning Molecular Representation in a Cell (2024). URL http://arxiv.org/abs/2406.12056. 2406.12056 [cs].

[35] Verleysen, M. & Fran¾cois, D. Cabestany, J., Prieto, A. & Sandoval, F. (eds) The Curse of Dimensionality in Data Mining and Time Series Prediction. (eds Cabestany, J., Prieto, A. & Sandoval, F.) Computational Intelligence and Bioinspired Systems, 758–770 (Springer, Berlin, Heidelberg, 2005).

[36] Radford, A. et al. Learning Transferable Visual Models From Natural Language Supervision (2021). URL http://arxiv.org/abs/2103.00020. 2103.00020

[37] Khosla, P. et al. Supervised Contrastive Learning (2021). URL http://arxiv.org/abs/2004.11362. 2004.11362 [cs].

[38] Wei, J. & Zou, K. EDA: Easy Data Augmentation Techniques for Boosting Performance on Text Classification Tasks (2019). URL http://arxiv.org/abs/1901.11196. 1901.11196 [cs].

[39] Schuh, M. G., Boldini, D. & Sieber, S. A. Synergizing Chemical Structures and Bioassay Descriptions for Enhanced Molecular Property Prediction in Drug Discovery. Journal of Chemical Information and Modeling 64, 4640–4650 (2024). URL 10.1021/acs.jcim.4c00765.

[40] Devlin, J., Chang, M.-W., Lee, K. & Toutanova, K. BERT: Pre-training of Deep Bidirectional Transformers for Language Understanding (2019). URL http://arxiv.org/abs/1810.04805. 1810.04805 [cs].

[41] Lee, J. et al. BioBERT: a pre-trained biomedical language representation model for biomedical text mining. Bioinformatics 36, 1234–1240 (2020). URL 10.1093/bioinformatics/btz682.

[42] Gorishniy, Y., Kotelnikov, A. & Babenko, A. TabM: Advancing Tabular Deep Learning with Parameter-Efficient Ensembling (2025). URL http://arxiv.org/abs/2410.24210. 2410.24210 [cs].

[43] Szklarczyk, D. et al. The STRING database in 2023: protein–protein association networks and functional enrichment analyses for any sequenced genome of interest. Nucleic Acids Research 51, D638–D646 (2023). URL https://doi.org/10.1093/nar/gkac1000.

[44] Cacace, E. et al. Uncovering nitroxoline activity spectrum, mode of action and resistance across Gram-negative bacteria. Nature Communications 16, 3783 (2025). URL https://www.nature.com/articles/s41467-025-58730-5. Publisher: Nature Publishing Group.

[45] Seyfert, C. E. et al. New Genetically Engineered Derivatives of Antibacterial Darobactins Underpin Their Potential for Antibiotic Development. Journal of Medicinal Chemistry 66, 16330–16341 (2023). URL 10.1021/acs.jmedchem.3c01660.

[46] Imai, Y. et al. A new antibiotic selectively kills Gram-negative pathogens. Nature 576, 459–464 (2019). URL https://www.nature.com/articles/s41586-019-1791-1. Publisher: Nature Publishing Group.

[47] Kany, A. M. et al. In Vivo Activity Profiling of Biosynthetic Darobactin D22 against Critical Gram-Negative Pathogens. ACS Infectious Diseases 10, 4337– 4346 (2024). URL 10.1021/acsinfecdis.4c00687.

[48] Xiao, Y., Gerth, K., Müller, R. & Wall, D. Myxobacterium-Produced Antibiotic TA (Myxovirescin) Inhibits Type II Signal Peptidase. Antimicrobial Agents and Chemotherapy 56, 2014–2021 (2012). URL https://journals.asm.org/doi/10. 1128/aac.06148-11. Publisher: American Society for Microbiology.

[49] Hübner, I. et al. Broad Spectrum Antibiotic Xanthocillin X Effectively Kills Acinetobacter baumannii via Dysregulation of Heme Biosynthesis. ACS Central Science 7, 488–498 (2021). URL https://pubs.acs.org/doi/10.1021/acscentsci.0c01621.

[50] Testolin, G. et al. Synthetic studies of cystobactamids as antibiotics and bacterial imaging carriers lead to compounds with high in vivo efficacy. Chemical Science 11, 1316–1334 (2020). URL https://pubs.rsc.org/en/content/articlelanding/2020/sc/c9sc04769g.

[51] Baumann, S. et al. Cystobactamids: Myxobacterial Topoisomerase Inhibitors Exhibiting Potent Antibacterial Activity. Angewandte Chemie International Edition 53, 14605–14609 (2014). URL https://onlinelibrary.wiley.com/doi/abs/10.1002/anie.201409964. eprint: https://onlinelibrary.wiley.com/doi/pdf/10.1002/anie.201409964.

[52] Kohnhöuser, D. et al. Optimization of the Central α-Amino Acid in Cystobactamids to the Broad-Spectrum, Resistance-Breaking Antibiotic CN-CC-861. Journal of Medicinal Chemistry 67, 17162–17190 (2024). URL https://doi.org/10.1021/acs.jmedchem.4c00927. Publisher: American Chemical Society.

[53] Subanovic, M., Frawley, D., Tierney, C., Velasco-Torrijos, T. & Walsh, F. Proteomic and metabolomic responses of priority bacterial pathogens to subinhibitory concentration of antibiotics. npj Antimicrobials and Resistance 3, 80 (2025). URL https://www.nature.com/articles/s44259-025-00147-7. Publisher: Nature Publishing Group.

[54] Jiang, J. et al. Boosting Tree-Assisted Multitask Deep Learning for Small Scientific Datasets. Journal of chemical information and modeling 60, 1235–1244 (2020). URL https://pmc.ncbi.nlm.nih.gov/articles/PMC7350172/.

[55] Köllen, M. F. et al. Generative Deep Learning Pipeline Yields Potent Gram-Negative Antibiotics. JACS Au 5, 4249–4259 (2025). URL 10.1021/jacsau.5c00602.

[56] Coscia, F. et al. A streamlined mass spectrometry–based proteomics workflow for large-scale FFPE tissue analysis. The Journal of Pathology 251, 100–112 (2020). URL https://onlinelibrary.wiley.com/doi/abs/10.1002/path.5420. eprint: https://pathsocjournals.onlinelibrary.wiley.com/doi/pdf/10.1002/path.5420.

[57] Bottlinger, M. et al. Natural product-derived ianthelliformisamines inhibit protein translation and block bacterial flagellum assembly. ACS Chemical Biology (2026). Accepted for publication, February 25, 2026. Manuscript ID: cb-2025-010188.R2.

[58] Demichev, V., Messner, C. B., Vernardis, S. I., Lilley, K. S. & Ralser, M. DIA-NN: neural networks and interference correction enable deep proteome coverage in high throughput. Nature Methods 17, 41–44 (2020). URL https://www.nature.com/articles/s41592-019-0638-x.

[59] McInnes, L., Healy, J. & Melville, J. UMAP: Uniform Manifold Approximation and Projection for Dimension Reduction (2020). URL http://arxiv.org/abs/1802.03426. 1802.03426 [stat].

[60] Landrum, G. et al. rdkit/rdkit: Release 2023.09.5 (2024). URL https://doi.org/10.5281/zenodo.10633624.

[61] OpenAI. ChatGPT. https://chatgpt.com/ (2025). Used August 8, 2025.

[62] Paszke, A. et al. PyTorch: An Imperative Style, High-Performance Deep Learning Library (2019). URL http://arxiv.org/abs/1912.01703. 1912.01703 [cs].

[63] Pedregosa, F. et al. Scikit-learn: Machine Learning in Python (2018). URL http://arxiv.org/abs/1201.0490. 1201.0490 [cs].

[64] Chen, T. & Guestrin, C. Krishnapuram, B. et al. (eds) Xgboost: A scalable tree boosting system. (eds Krishnapuram, B. et al.) Proceedings of the 22nd ACM SIGKDD International Conference on Knowledge Discovery and Data Mining, KDD ‘16, 785–794 (Association for Computing Machinery, New York, NY, USA, 2016). URL 10.1145/2939672.2939785.

[65] Ke, G. et al. Guyon, I. et al. (eds) LightGBM: A Highly Efficient Gradient Boosting Decision Tree. (eds Guyon, I. et al.) Advances in Neural Information Processing Systems, Vol. 30 (Curran Associates, Inc., 2017). URL https://proceedings.neurips.cc/paperfiles/paper/2017/file/6449f44a102fde848669bdd9eb6b76fa-Paper.pdf.

[66] Ash, J. R. et al. Practically Significant Method Comparison Protocols for Machine Learning in Small Molecule Drug Discovery. Journal of Chemical Information and Modeling 65, 9398–9411 (2025). URL 10.1021/acs.jcim.5c01609.

[67] Hendrycks, D. & Gimpel, K. A Baseline for Detecting Misclassified and Out-of-Distribution Examples in Neural Networks (2018). URL http://arxiv.org/abs/1610.02136. 1610.02136 [cs].

[68] Houlsby, N., Huszár, F., Ghahramani, Z. & Lengyel, M. Bayesian Active Learning for Classification and Preference Learning (2011). URL http://arxiv.org/abs/1112.5745. 1112.5745 [stat].

[69] Gal, Y., Islam, R. & Ghahramani, Z. Precup, D. & Teh, Y. W. (eds) Deep Bayesian active learning with image data. (eds Precup, D. & Teh, Y.W.) Proceedings of the 34th International Conference on Machine Learning, Vol. 70 of Proceedings of Machine Learning Research, 1183–1192 (PMLR, 2017). URL https://proceedings.mlr.press/v70/gal17a.html.

[70] Lakshminarayanan, B., Pritzel, A. & Blundell, C. Simple and Scalable Predictive Uncertainty Estimation using Deep Ensembles (2017). URL http://arxiv.org/abs/1612.01474. 1612.01474 [stat].

[71] Perez-Riverol, Y. et al. The PRIDE database at 20 years: 2025 update. Nucleic Acids Research 53, D543–D553 (2025). URL 10.1093/nar/gkae1011.

