## Supplementary information for "Decoding antibiotic modes of action from multimodal cellular responses"

**3 Supplementary tables** **11**

### 1 Methods

#### 1.1 Overview

This Supplementary Information provides additional methodological detail for the multimodal mode of action (MoA) prediction framework, with emphasis on (i) the exact construction of feature matrices and the compound–MoA pairwise learning task, and (ii) the statistical analyses used to compare augmentation strategies, feature sets, model families and uncertainty estimators. Supplementary Tables S1 and S2 summarize compound sources and library composition; Supplementary Tables S3, S4, and S5 collect proteomics experimental details; and Supplementary Tables S6, S7, and S8 provide compact statistical summaries for Figures 3–5.

#### 1.2 Data objects and modality matrices

All modeling and figure-generation notebooks and scripts consume precomputed, sample-indexed modality matrices stored as serialized tables under `Data/Base/`. These include proteomics (`X_proteomics.pkl` and a significance-filtered variant `X_proteomics_sig.pkl`), chemical structure fingerprints (`X_ecfp.pkl`), InfoAlign embeddings (`X_cellpaint.pkl`), MIC features (`X_mic.pkl`), growth curve features (`X_gc.pkl`), and MoA labels (`y_labels.pkl`). Sample identifiers follow a compound-and-replicate convention (for example, `JH<compound>_<rep>`). Mapping of sample identifiers (`JH<compound>`) to antibiotics is shown in Supplementary Tables S3.

#### 1.3 Proteomics feature matrix construction

Gene-level proteomics matrices are assembled from Perseus export tables by retaining quantitative intensity columns and a gene-annotation column, expanding rows with multiple gene assignments, and building a unified gene-by-sample matrix over the *E. coli* K-12 UniProt FASTA gene universe. Missing values are filled with zeros and all values are coerced to numeric values.

For analyses requiring improved portability across mass spectrometry platforms or laboratories, a significance-filtered proteomics representation is used. In this representation, replicate intensities for proteins that do not meet a per-protein significance criterion relative to matched controls are set to zero. The external and Eclipse dataset processing notebooks implement this by thresholding $-\log_{10}(p)$ at 1.3 (approximately $p < 0.05$) and zeroing all replicate intensity values failing the threshold.

Table S5.

#### 1.4 Chemical structure features

Chemical structure features are computed from SMILES strings as Morgan fingerprints (ECFP-like) using RDKit with radius 2 and 2,048 bits.

#### 1.5 InfoAlign embeddings ("CellPaint" modality)

The auxiliary morphological–transcriptional modality is represented by 300-dimensional embeddings predicted from SMILES using a pretrained InfoAlign representer. In the codebase and figure panels, this modality is referred to as `CellPaint` but displayed as `InfoAlign`.

#### 1.6 MIC and growth curve features

MIC features are computed by mapping measured MIC values onto a two-fold dilution series (from 800 down to 0.001), filling unmeasured steps by a nearest-value rule, and applying per-compound min–max scaling to yield a 20-dimensional vector.

Growth curve features are computed by selecting, per compound, the lowest tested antibiotic concentration whose final-timepoint $OD_{600}$ (24 h) falls below 50% of the solvent control (or the highest tested concentration otherwise). The selected $OD_{600}$ time series is multiplied by the concentration factor for that condition and summarized as a 10 time-point vector.

#### 1.7 MoA text descriptions, augmentation and embeddings

For each MoA class, a mechanistic description is curated and expanded into ten text variants per augmentation regime. All generated outputs were manually reviewed, screened for factual consistency and redundancy, and edited where necessary before downstream analysis.

- Sentence augmentation shuffles the sentence order within a description.

- Word augmentation shuffles word order.
- Semantic augmentation uses paraphrases intended to preserve mechanistic meaning while varying wording; these paraphrases were generated with ChatGPT (OpenAI) [1].

Text embeddings are computed using transformer encoders (PubChemDeBERTa-augmented, BERT-base-uncased and BioBERT v1.1) [2–4]. Embeddings use mean pooling over last-layer hidden states and have dimensionality 768. In embedding generation, text is truncated to a fixed maximum length (128 tokens in the embedding helper).

###### 1.7.1 Exact text-encoder checkpoint revisions

For reproducibility, we report the exact Hugging Face Hub snapshot revisions (git commit SHAs) resolved for each text encoder during embedding generation:

- `mschuh/PubChemDeBERTa-augmented@955028022c9ae27879aeaa044ba4f870cef3b06b`
- `google-bert/bert-base-uncased@86b5e0934494bd15c9632b12f734a8a67f723594`
- `dmis-lab/biobert-v1.1@551ca18efd7f052c8dfa0b01c94c2a8e68bc5488`

##### 1.8 Pairwise learning task and multiclass readout

The primary learning formulation is a compound–MoA pairwise binary classification task. Each experimental sample is crossed with each candidate MoA embedding (and its augmentation index), producing rows that combine selected sample modalities with the candidate text embedding. The binary target is defined as 1 if the candidate MoA equals the true MoA label and 0 otherwise. A default probability threshold of 0.5 is used when converting binary scores to class labels.

For multiclass summaries derived from pairwise scores, pairwise scores are pivoted to a sample-by-candidate matrix and averaged across text augmentations before selecting the highest-scoring MoA (argmax). No-text baselines use sample-level feature matrices without pairwise expansion.

##### 1.9 Models and hyperparameters

###### 1.9.1 TabM

The primary deployment model is TabM [5]. The wrapper trains for up to 100 epochs with batch size 256, learning rate $10^{-4}$ and weight decay $3 \times 10^{-4}$, with early stopping (patience 3). The TabM implementation uses an internal `QuantileTransformer` (output distribution "normal") fit on the training data.

###### 1.9.2 Neural and classical baselines

The augmentation benchmark uses a PyTorch Lightning multilayer perceptron [6]. The broader model screen additionally includes a scikit-learn multilayer perceptron [7], LightGBM[8], XGBoost [9] and a scikit-learn random forest [7]. These baselines are run under the same compound-level split protocol and evaluated using identical metrics.

###### 1.9.3 Baseline model configurations (non-TabM)

Unless stated otherwise, baseline model hyperparameters correspond to the factory defaults used in the evaluation scripts.

**PyTorch MLP (Lightning; `pytorch_mlp`).** The Lightning MLP wrapper is configured with `batch_size=128`, `max_epochs=40`, `width_list=[512,256]`, `depth=2`, `activation="relu"`, `dropout=0.5`, and `lr=1e-4`. Each hidden block applies a linear layer, batch normalization (`BatchNorm1d`), dropout and activation. Weights are initialized with Xavier uniform initialization. Training uses Adam with `weight_decay=1e-4` and early stopping on epoch-averaged `train_loss` with `patience=5`. For binary tasks, the loss is `BCEWithLogitsLoss` and probabilities are obtained with a sigmoid; for multiclass tasks, the loss is `CrossEntropyLoss` and probabilities are obtained with a softmax. Binary class predictions use a threshold of 0.5 on the positive-class probability.

**scikit-learn MLP (`mlp`).** The scikit-learn multilayer perceptron is `MLPClassifier(hidden_layer_sizes=(256,128), max_iter=400)` with `random_state` set by the run seed.

**LightGBM (`lightgbm`).** LightGBM uses `LGBMClassifier(n_estimators=200, random_state=seed)` with `subsample` implemented via `bagging_fraction=0.9` and `bagging_freq=1`, and feature subsampling via `feature_fraction=0.9`. Additional settings include `min_data_in_leaf=1`, `min_data_in_bin=1`, `feature_pre_filter=False`, and `verbosity=-1`. For binary tasks, `class_weight="balanced"` is used; for multiclass tasks, `objective="multiclass"` and `num_class=n_classes` are set.

**XGBoost (`xgboost`).** XGBoost uses `XGBClassifier(n_estimators=300, subsample=0.9, colsample_bytree=0.9, random_state=seed, use_label_encoder=False)`. For binary tasks, `eval_metric="logloss"` is used; for multiclass tasks, `objective="multi:softprob"`, `num_class=n_classes`, and `eval_metric="mlogloss"` are set.

**Random forest (`rf`).** Random forests use `RandomForestClassifier(n_estimators=400, random_state=seed)`. For binary tasks, `class_weight="balanced"` is used; for multiclass tasks, no class weighting is applied.

##### 1.10 Cross-validation protocols and leakage control

All evaluation splits are performed at the compound level. In repeated stratified $K$-fold cross-validation, compounds are stratified by their MoA label and assigned to folds; all pairwise-expanded rows inherit the fold assignment of their parent compound. Repeated cross-validation uses a deterministic seed schedule with fold random state `base_seed + 1000`×`iteration`, and model replica seeds `base_seed + rep_idx`.

Uncertainty estimation experiments use leave-one-out (LOO; held-out compound) and leave-one-class-out (LOCO; held-out MoA class) protocols with multiple seed repeats.

##### 1.11 Evaluation metrics

Binary performance metrics include PR AUC, ROC AUC and Matthews correlation coefficient (MCC). Multiclass summaries reported in this manuscript include macro-F1 and one-vs-rest macro PR AUC and macro ROC AUC. Top-1 class selection is used to derive final predicted MoA labels from aggregated class scores, and top-3 class displays are used descriptively in selected prediction plots.

#### 1.12 Uncertainty meta-model and calibration

The deployed uncertainty estimator is a logistic-regression meta-model trained on derived uncertainty features from the MoA prediction outputs. The model is implemented as:

$$\texttt{StandardScaler} \rightarrow \texttt{PolynomialFeatures}(\text{degree} = 2)$$
$$\rightarrow \texttt{LogisticRegression}(\text{class\_weight=balanced})$$

and is calibrated using `CalibratedClassifierCV` with sigmoid calibration and three-fold internal cross-validation. During external application, floor-based overrides are applied before final thresholding to conservatively label low-signal cases as uncertain.

#### 1.13 Derived uncertainty features

Let $\mathbf{p}_i^{(u)} \in [0,1]^C$ denote the perturbation-level multiclass distribution for compound $i$ and perturbation index $u = 1, \ldots, M_i$, where perturbations arise from augmentation, seed, and expert variation after pairwise scores have been aggregated to the multiclass level. The perturbation-averaged class distribution is

$$\bar{\mathbf{p}}_i = \frac{1}{M_i} \sum_{u=1}^{M_i} \mathbf{p}_i^{(u)}.$$

From this distribution we define the compound-level top score and margin as

$$\text{PredScore}_i = \max_c \bar{p}_{i,c}, \qquad \text{Margin}_i = \bar{p}_{i,c_i^*} - \bar{p}_{i,c_i^{(2)}},$$

where $c_i^* = \arg\max_c \bar{p}_{i,c}$ and $c_i^{(2)}$ is the runner-up class. The entropy-based features are

$$\text{PredEntropy}_i = H(\bar{\mathbf{p}}_i), \qquad \text{ExpEntropy}_i = \frac{1}{M_i} \sum_{u=1}^{M_i} H(\mathbf{p}_i^{(u)}),$$

with $H(\mathbf{p}) = -\sum_c p_c \log p_c$, and

$$\text{EpistemicMI}_i = \text{PredEntropy}_i - \text{ExpEntropy}_i.$$

Disagreement features were computed as

$$\text{MeanPairKL}_i = \frac{1}{|\mathcal{P}_i|} \sum_{(u,v) \in \mathcal{P}_i} \text{KL}\left(\mathbf{p}_i^{(u)} \middle\| \mathbf{p}_i^{(v)}\right),$$

where $\mathcal{P}_i$ is the set of sampled perturbation pairs used in the implementation,

$$\text{KConsistency}_i = \max_c \frac{1}{M_i} \sum_{u=1}^{M_i} \mathbf{1}\left[\arg\max_k p_{i,k}^{(u)} = c\right], \qquad \text{VariationRatio}_i = 1 - \text{KConsistency}_i,$$

and

$$\text{VarMax}_i = \text{Var}_u\left(\max_c p_{i,c}^{(u)}\right).$$

For class-specific stability features, let $y_i^* = \arg\max_c \bar{p}_{i,c}$ be the predicted compound-level class. Let $s_{i,z}$ denote the predicted-class score for seed $z$, and $s_{i,z,a}$ the corresponding score for seed $z$ and augmentation $a$. We define

$$\text{SeedStd}_i = \text{sd}_z(s_{i,z}), \qquad \text{AugVar}_i = \frac{1}{Z_i} \sum_z \text{Var}_a(s_{i,z,a}).$$

For biological replicates, let $s_{i,r}^{\text{bio}}$ denote the score of the predicted class $y_i^*$ in replicate $r$, and let

$$m_{i,r}^{\text{rep}} = p_{i,c_r^*}^{(r)} - p_{i,c_r^{(2)}}^{(r)}$$

be the replicate-level top-vs-second margin, where $c_r^*$ and $c_r^{(2)}$ are the top and runner-up classes within replicate $r$. Then

$$\text{BioVar}_i = \text{Var}_r(s_{i,r}^{\text{bio}}), \qquad \text{RepMarginStd}_i = \text{sd}_r(m_{i,r}^{\text{rep}}),$$

$$\text{RepMarginIQR}_i = Q_{0.75}(m_{i,r}^{\text{rep}}) - Q_{0.25}(m_{i,r}^{\text{rep}}), \qquad \text{RepAgree}_i = \frac{1}{R_i}\sum_{r=1}^{R_i} \mathbf{1}[c_r^* = y_i^*]\,.$$

These features form the candidate input space for the logistic-regression uncertainty meta-model; for transfer inference, the fitted error probability is additionally subjected to the floor-based overrides described in the main Methods.

##### 1.14 Reproducibility notes

Randomness controls are applied consistently across data splits, model training and text-embedding generation (explicit seeding for Python, NumPy and PyTorch; deterministic transformer execution in the embedding helper). Run-level provenance is captured via serialized metadata (for example, `meta.json` and `predict_manifest.json`).

#### 2 Supplementary figures and statistical analyses

This section groups all supplementary figures together with the statistical details and machine-readable summaries that support them.

##### 2.1 Reporting conventions

Unless otherwise stated, inferential tests are two-sided with $\alpha = 0.05$. Exact $p$-values are reported. Multiple comparisons are controlled using Bonferroni adjustment for figure-level omnibus testing across metrics, Tukey HSD for ANOVA post hoc comparisons, Benjamini–Hochberg (BH) correction for Conover–Friedman post hoc comparisons, and Holm adjustment for uncertainty-method pairwise tests, as indicated per analysis. Effect sizes in uncertainty-method comparisons are reported as Cliff's delta with bootstrap confidence intervals ($N = 5000$ resamples).

#### 2.2 Figure 3

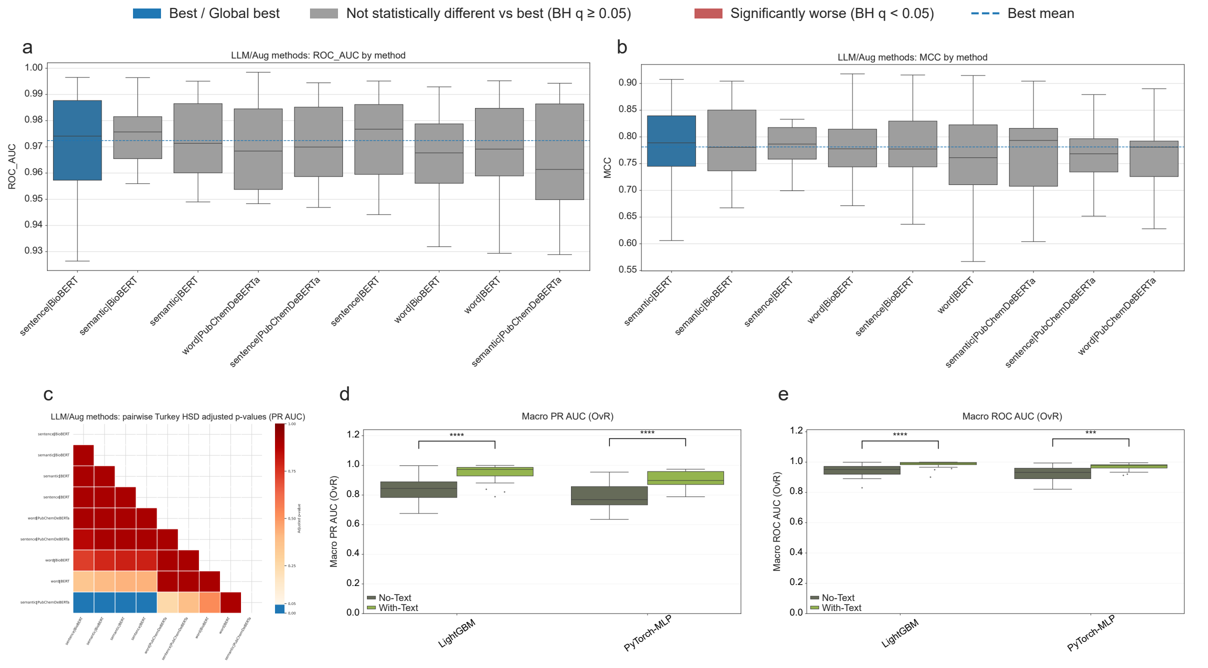

Figure S1: **Extended benchmarking and statistical analysis for Fig. 3**: (a,b) Additional LLM/augmentation benchmarking panels corresponding to Fig. 3c (ROC AUC and MCC), shown across 25 balanced cross-validation cycles (5 repeats $\times$ 5 folds) for the 9 augmentation/encoder combinations (PyTorch-MLP; Text Embedding + Proteomics). Boxplots show median (center line), interquartile range (box), whiskers (1.5$\times$IQR), and outliers (points). (c) PR AUC all-vs-all Tukey HSD-adjusted pairwise comparison matrix for the 9 augmentation/encoder methods. (d,e) Additional text-vs-no-text panels for Fig. 3d (macro PR AUC OvR and macro ROC AUC OvR), grouped by model (LightGBM, PyTorch-MLP); brackets indicate paired within-model comparisons using Bonferroni-corrected $t$-tests. Exact omnibus and post hoc test values are reported below and in Table S6.

- **Figure 3c** (9 augmentation/encoder methods; PyTorch-MLP with Text Embedding + Proteomics): inferential testing follows the repeated-measures workflow of Ash et al[10]. RM-ANOVA was run separately for PR AUC, ROC AUC and MCC across 25 cross-validation cycles, then Bonferroni correction was applied across these three omnibus tests ($\alpha_{\text{Bonf}} = 0.05/3 = 0.0167$)(Fig. S1 a-b). Observed omnibus $p$-values were PR AUC $= 1.96 \times 10^{-3}$, ROC AUC $= 2.00 \times 10^{-2}$ and MCC $= 1.75 \times 10^{-2}$; therefore only PR AUC passed the Bonferroni-gated omnibus criterion.

- **Figure 3c post hoc and coloring rule:** Tukey HSD pairwise comparisons are performed only for metrics passing the Bonferroni-gated omnibus test. For PR AUC, Tukey-adjusted pairwise $p$-values were computed for all method pairs; the full corrected p-value heatmap is shown in Fig. S1 c). For coloring of the main-figure boxplots, this full pairwise matrix was reduced to comparisons against the best-mean method (`sentence|BioBERT`, mean PR AUC $= 0.8869$), with blue = best, gray = not different from best, and red = significantly worse.

- **Figure 3d** (4 groups: LightGBM/PyTorch-MLP $\times$ with-text/no-text): Friedman omnibus tests were computed per metric across 25 balanced cross-validation cycles. Bonferroni correction was applied across the three figure-level metrics using adjusted alpha only ($\alpha_{\mathrm{Bonf}} = 0.05/3 = 0.0167$; decision rule $p_{\mathrm{raw}} < \alpha_{\mathrm{Bonf}}$). All three metrics passed this threshold (macro-F1: $p = 1.04 \times 10^{-6}$; macro PR AUC OvR: $p = 1.31 \times 10^{-10}$; macro ROC AUC OvR: $p = 5.01 \times 10^{-11}$). Within-model paired $t$-tests (with-text minus no-text), Bonferroni-corrected across the three metrics per model, were significant in all six cases (Table S6).

#### 2.3 Figure 4

- **Figure 4a** (feature-set comparisons): inferential testing was performed on balanced cross-validation cycles using Friedman omnibus tests (within-block repeated design; full ROC AUC and MCC feature-set panels in Fig. S2a,b). We used Friedman rather than RM-ANOVA because this benchmark compares many repeated conditions (32 feature coalitions per model) and bounded performance metrics, where normality/sphericity assumptions are less reliable and a rank-based repeated-block test is more robust. In the full model-by-metric analysis (6 models $\times$ 3 metrics, 32 feature sets), Bonferroni correction was applied across the three metrics within each model ($\alpha_{\mathrm{Bonf}} = 0.05/3 = 0.0167$; decision rule $p_{\mathrm{raw}} < \alpha_{\mathrm{Bonf}}$). All omnibus tests were highly significant (raw $p$ values in the range $\sim 10^{-56}$ to $\sim 10^{-84}$), so post hoc testing remained applicable for all model-metric combinations. Top-3 PR AUC coalitions per model are summarized in Table S7.

- **Figure 4a post hoc**: pairwise feature-set comparisons used Conover–Friedman with Benjamini–Hochberg correction to compute a full all-vs-all adjusted $q$-value matrix (visualized as pairwise heatmaps in Fig. S2c–e). For interpretation in the boxplot coloring and summary tables, this full matrix was then reduced to a best-vs-rest view: for each model-metric combination, the best-mean coalition was identified, all other coalitions were compared against this reference, and coalitions with BH-adjusted $q \geq 0.05$ were reported as the *not-significantly-worse-than-best* set.

- **Figure 4b** (global comparison across 18 conditions = 6 models $\times$ 3 selected feature sets): Friedman test over 25 balanced cross-validation cycles yielded $Q = 148.16$, $df = 17$, $p = 5.61 \times 10^{-23}$; post hoc method was Conover–Friedman (BH) versus the best condition. Companion ROC AUC and MCC boxplot panels are shown in Fig. S2f,g, and the corresponding pairwise BH-adjusted $q$-value heatmaps are shown in Fig. S2i,j (with PR AUC heatmap in Fig. S2h).

- **Figure 4c** (exact Shapley values) and **Figure 4d–h** (compound-level probability profiles) are descriptive in this manuscript version; no additional inferential tests were applied to these panels.

Complete per-cycle scores, omnibus test outputs, pairwise BH-adjusted matrices, and full coalition rankings for all three metrics (visualized in Fig. S2) are provided in the machine-readable exports (`analysis_exports/Feature_augmentation_experiment/`).

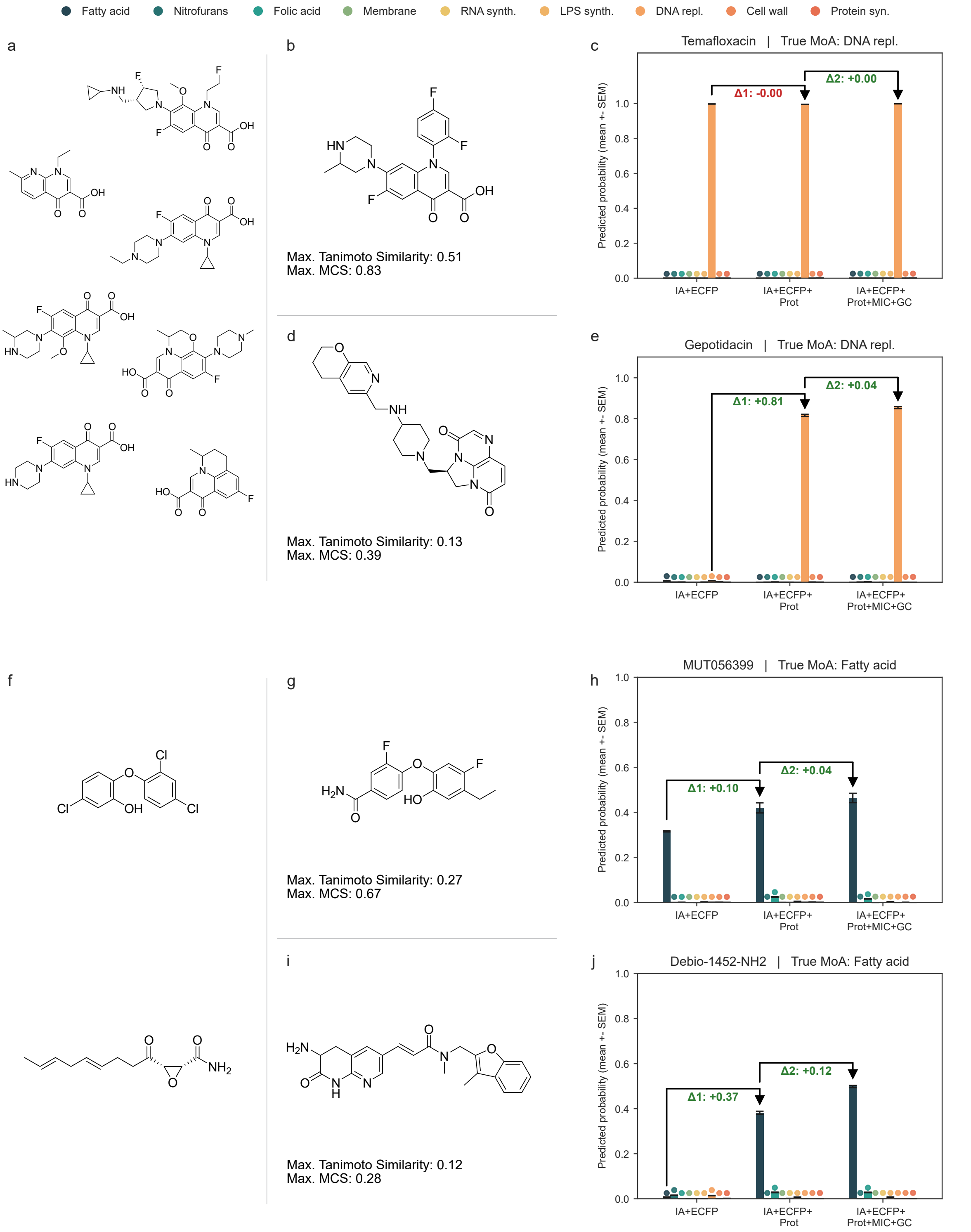

Figure S2: **Extended feature-set benchmarking and statistical analysis for Fig. 4**: (a,b) TabM performance across all evaluated feature coalitions for ROC AUC (a) and MCC (b). Boxplots show median (center line), IQR (box), whiskers (1.5×IQR), and outliers (points). (c–e) All-vs-all Conover–Friedman pairwise comparison matrices (BH-adjusted $q$-values) for TabM on PR AUC (c), ROC AUC (d), and MCC (e). (f,g) Cross-model comparison for the three selected high-performing coalitions for ROC AUC (f) and MCC (g), with global Friedman results shown in panel titles. (h–j) All-vs-all pairwise BH-adjusted $q$-value matrices for the model–coalition combinations in (f,g), shown for PR AUC (h), ROC AUC (i), and MCC (j).

#### 2.4 Figure 5

All methods were calibrated to a fixed in-distribution false-positive-rate target (target_id_fpr=0.10). Overall benchmark performance across the six uncertainty methods is summarized in Table S8. The benchmark dataset contains 51 compounds. For inferential method-comparison on the primary metric (err_recall), analysis used 39 compounds across six uncertainty methods: 12 compounds had n_err_sum=0 (no observed errors across the aggregated LOO+LOCO evaluation), making per-compound err_recall undefined (NaN), and were therefore excluded from the inferential table used for omnibus/pairwise testing.

- Omnibus test: RM-ANOVA statistic = 13.74, $p = 1.86 \times 10^{-11}$.
- Pairwise tests: Holm-adjusted multiple comparisons across all 15 method pairs; three pairs reached significance at 5% (High expected entropy vs. Meta-model (LR+Poly), $p_{\text{Holm}} = 1.81 \times 10^{-2}$; Low max score vs. Z-Gate (global), $p_{\text{Holm}} = 4.79 \times 10^{-2}$; Meta-model (LR+Poly) vs. Z-Gate (global), $p_{\text{Holm}} = 1.35 \times 10^{-2}$).
- Corresponding Cliff's delta values were 0.538, −0.513, and −0.590, respectively.

#### 2.5 Figure 6

Figure 6 was treated as descriptive out-of-distribution validation (small-$n$ panels); no inferential hypothesis tests were run. Uncertainty calls used the recalibrated artifact models/uncertainty_estimator_adapted/meta_artifact.json (meta_threshold=0.2779), where recalibration used only development LOO/LOCO semantic+proteomics prediction metadata (no external or Eclipse test labels). Under these settings, all compounds were flagged uncertain in the unfiltered Eclipse and external-data panels, all three Eclipse compounds were confidently classified after significance filtering, and only Imipenem remained confident in the significance-filtered Subanovic panel.

#### 2.6 Machine-readable statistical tables

All compact CSV summaries used for this Supplementary Information section are exported under analysis_exports/supplementary_stats/, including figure-level omnibus tests, pairwise test outputs, and panel-level uncertainty summaries. A dedicated regeneration notebook is provided at notebooks/supplementary/regenerate_supplementary_stats.ipynb, which rebuilds these manuscript-facing CSVs from the primary figure-analysis exports and prediction outputs in this repository.

### 3 Supplementary tables

This section collects all supplementary tables referenced in the manuscript and Methods, including compound provenance, library composition, proteomics acquisition details, and compact statistical summaries for Figures 3–5.

Table S1: Vendor source information for all compounds.

| **Compound** | **Source (company, catalog no.)** |
|---|---|
| Ampicillin | Karl Roth, KB029 |
| Amoxicillin | MedChemExpress, HY-B0467A |
| Sulopenem | MedChemExpress, HY-105284 |
| Cefodizime | MedChemExpress, HY-108402 |
| Aztreonam | MP Biomedicals, 150415 |
| Doripenem | MedChemExpress, HY-B0187 |
| Imipenem (monohydrate) | MedChemExpress, HY-B1369 |
| Ertapenem (disodium) | MedChemExpress, HY-A0294A |
| Cefiderocol | MedChemExpress, HY-17628 |
| Cefepime | MedChemExpress, HY-B0692 |
| Fosfomycin (calcium) | MedChemExpress, HY-B1075 |
| Retapamulin | MedChemExpress, HY-17010 |
| Framycetin (sulfate) | MedChemExpress, HY-17624A |
| Chloramphenicol | Karl Roth, 3886 |
| Tigecycline | MedChemExpress, HY-B0117 |
| Tetracycline | MedChemExpress, HY-A0107 |
| Azithromycin | MedChemExpress, HY-17506 |
| Gentamicin sulfate | Karl Roth, 0233 |
| Eravacycline (dihydrochloride) | MedChemExpress, HY-16980A |
| Apramycin | MedChemExpress, HY-B1329 |
| Kanamycin sulfate | Karl Roth, T832 |
| Tobramycin | Acros Organics, 455430010 |
| Sulfamonomethoxine | MedChemExpress, HY-B0946 |
| Sulfisoxazole | MedChemExpress, HY-B0323 |
| Sulfacetamide | MedChemExpress, HY-N7123 |
| Sulfaguanidine | MedChemExpress, HY-B1267 |
| Sulfadiazine | Thermo Scientific, A12370.18 |
| Sulphanilamide | Karl Roth, 4716 |
| Trimethoprim | Acros Organics, 455120050 |
| Tetroxoprim | MedChemExpress, HY-107033 |
| Gatifloxacin | MedChemExpress, HY-10581 |
| Temafloxacin | MedChemExpress, HY-16487 |
| Lascufloxacin | MedChemExpress, HY-16745 |
| Ciprofloxacin | Sigma, 17850 |
| Levofloxacin | Alfa Aesar, J66943 |
| Gepotidacin | MedChemExpress, HY-16742 |
| Nalidixic acid | MedChemExpress, HY-B0398 |
| Novobiocin | MedChemExpress, HY-B0425 |
| Flumequine | MedChemExpress, HY-B0526 |
| Enrofloxacin | Sigma, 17849 |
| Nitrofurantoin | Sigma, N7878 |
| Nitrofurazone | TCI, N0200 |
| D12 | Synthesized in-house according to Köllen *et al.*[11] |
| D8 | Synthesized in-house according to Köllen *et al.*[11] |
| Rifaximin | MedChemExpress, HY-13234 |
| Rifampicin | Karl Roth, 4163 |
| Rifabutin | MedChemExpress, HY-17025 |
| Triclosan | Karl Roth, 1KNA |
| Cerulenin | MedChemExpress, HY-A0210 |
| MUT056399 | MedChemExpress, HY-18169 |
| Debio-1452-$NH_2$ | Synthesized in-house according to Parker *et al.*[12] |
| Colistin (sulfate) | MedChemExpress, HY-A0089 |
| Polymyxin B | MedChemExpress, HY-149179 |
| Squalamine | MedChemExpress, HY-16468 |
| Thanatin (TFA) | MedChemExpress, HY-P5601A |
| Lolamicin | MedChemExpress, HY-164036 |
| PF-5081090 | MedChemExpress, HY-103251 |
| ACHN-975 (TFA) | MedChemExpress, HY-19936A |
| CHIR-090 | MedChemExpress, HY-15460 |
| BB-78485 | Synthesized in-house by adapting Shao *et al.*[13] |
| Xanthocillin | Synthesized in-house according to Hübner *et al.*[14] |
| Myxovirescin | Provided by Helmholtz Institute for Pharmaceutical Research Saarland |
| Darobactin D22 | Provided by Helmholtz Institute for Pharmaceutical Research Saarland |
| Cystobactamid CN-DM-861 | Provided by Helmholtz Institute for Pharmaceutical Research Saarland |
| Nitroxoline | TCI, H0805 |
| Clioquinol | TCI, C0187 |
| Luteolin | Karl Roth, 4546 |
| Actinonin | Sigma, A6671 |

Table S2: **Antibiotics library and its usage in model development (base) or validation**.

| Compound | MIC | Mode of Action | Usage (Base / Validation / not used) |
|---|---|---|---|
| Ampicillin | 6.25 | Cell Wall Biosynthesis | Base |
| Amoxicillin | 50.00 | Cell Wall Biosynthesis | Not used |
| Sulopenem | 0.06 | Cell Wall Biosynthesis | Base |
| Cefodizime | 0.39 | Cell Wall Biosynthesis | Base |
| Aztreonam | 0.63 | Cell Wall Biosynthesis | Not used |
| Doripenem | 0.25 | Cell Wall Biosynthesis | Base |
| Imipenem | 0.31 | Cell Wall Biosynthesis | Base |
| Ertapenem | 0.5 | Cell Wall Biosynthesis | Base |
| Cefiderocol | 0.63 | Cell Wall Biosynthesis | Base |
| Cefepime | 0.25 | Cell Wall Biosynthesis | Base |
| Fosfomycin | 50.00 | Cell Wall Biosynthesis | Base |
| Retapamulin | 12.50 | Protein Biosynthesis | Not used |
| Framycetin | 100.00 | Protein Biosynthesis | Base |
| Chloramphenicol | 12.50 | Protein Biosynthesis | Base |
| Tigecycline | 0.63 | Protein Biosynthesis | Base |
| Tetracycline | 1.25 | Protein Biosynthesis | Base |
| Azithromycin | 1.56 | Protein Biosynthesis | Base |
| Gentamicin | 0.39 | Protein Biosynthesis | Base |
| Eravacycline | 0.04 | Protein Biosynthesis | Base |
| Apramycin | $\gg 200$ | Protein Biosynthesis | Not used |
| Kanamycin | 1.56 | Protein Biosynthesis | Base |
| Tobramycin | 1.56 | Protein Biosynthesis | Base |
| Sulfamonomethoxine | 100.00 | Folic Acid Metabolism | Base |
| Sulfisoxazole | 200 | Folic Acid Metabolism | Base |
| Sulfacetamide | $\gg 200$ | Folic Acid Metabolism | Not used |
| Sulfaguanidine | $\gg 200$ | Folic Acid Metabolism | Not used |
| Sulfadiazine | $\gg 200$ | Folic Acid Metabolism | Not used |
| Sulphanilamide | $\gg 200$ | Folic Acid Metabolism | Not used |
| Trimethoprim | 1.25 | Folic Acid Metabolism | Base |
| Tetroxoprim | 12.50 | Folic Acid Metabolism | Base |
| Gatifloxacin | 0.06 | DNA Synthesis | Base |
| Temafloxacin | 0.16 | DNA Synthesis | Base |
| Lascufloxacin | 0.08 | DNA Synthesis | Base |
| Ciprofloxacin | 0.03 | DNA Synthesis | Base |
| Levofloxacin | 0.06 | DNA Synthesis | Base |
| Gepotidacin | 2.50 | DNA Synthesis | Base |
| Nalidixic acid | 50.00 | DNA Synthesis | Base |
| Novobiocin | $\gg 200$ | DNA Synthesis | Not used |
| Flumequine | 2.50 | DNA Synthesis | Base |
| Enrofloxacin | 0.08 | DNA Synthesis | Base |
| Nitrofurantoin | 50.00 | Nitrofurans | Base |
| Nitrofurazone | 25.00 | Nitrofurans | Base |
| D12 | 50.00 | Nitrofurans | Base |
| D8 | 12.50 | Nitrofurans | Base |
| Rifaximin | 25.00 | RNA Synthesis | Base |
| Rifampicin | 12.50 | RNA Synthesis | Base |
| Rifabutin | 6.25 | RNA Synthesis | Base |
| Triclosan | 0.31 | Fatty Acid Synthesis | Base |
| Cerulenin | 400.00 | Fatty Acid Synthesis | Base |
| MUT056399 | 1.56 | Fatty Acid Synthesis | Base |
| Debio-1452-$NH_2$ | 6.25 | Fatty Acid Synthesis | Base |
| Colistin | 0.31 | Membrane Disruption | Base |
| Polymyxin B | 0.08 | Membrane Disruption | Base |
| Squalamine | 3.13 | Membrane Disruption | Base |
| Thanatin | 0.63 | Membrane Disruption | Base |
| Lolamicin | 2.50 | Membrane Disruption | Base |
| PF-5081090 | 0.31 | Lipopolysaccharide Synthesis | Base |
| ACHN-975 | 0.16 | Lipopolysaccharide Synthesis | Base |
| CHIR-090 | 0.31 | Lipopolysaccharide Synthesis | Base |
| BB-78485 | 1.56 | Lipopolysaccharide Synthesis | Base |
| Xanthocillin | 1.56 | Novel MoA | Validation |
| Cystobactamid CN-DM-861 | 0.31 | DNA Synthesis | Validation |
| Darobactin D22 | 0.78 | Novel MoA | Validation |
| Myxovirescin | 1.56 | Novel MoA | Validation |
| Nitroxoline | 12.50 | Novel MoA | Validation |
| Clioquinol | $\gg 200$ | Novel MoA | Not used |
| Luteolin | $\gg 200$ | Novel MoA | Not used |
| Actinonin | $\gg 200$ | Novel MoA | Not used |

Table S3: Compound-specific treatment conditions used for in-house proteomics sample generation. The table lists the internal compound identifiers, applied concentrations, MIC factors, and matched solvent controls for each proteomics experiment. Compounds marked with † correspond to proteomics measurements generated in the present project that were also included in Köllen *et al.*[11] and published there. Entries marked with ∗ indicate the corresponding sample identifiers used in the PRIDE upload for Köllen *et al.*

| **Compound identifier** | **Compound** | **Concentration for proteomics [$\mu$M]** | **MIC factor** | **Control** |
|---|---|---|---|---|
| JH0 / Amp* | Ampicillin† | 31.25 | 5×MIC | DMSO |
| JH2 | Sulopenem | 0.6 | 10×MIC | DMSO |
| JH3 | Cefodizime | 5.86 | 15×MIC | DMSO |
| JH5 | Doripenem | 2.5 | 10×MIC | DMSO |
| JH6 | Imipenem | 1.56 | 5×MIC | $H_2O$ |
| JH7 | Ertapenem | 2.50 | 5×MIC | DMSO |
| JH8 | Cefiderocol | 9.38 | 15×MIC | DMSO |
| JH9 | Cefepime | 1.25 | 5×MIC | DMSO |
| JH10 | Fosfomycin | 5 | 0.1×MIC | $H_2O$ + HCL (pH 2) |
| JH12 | Framycetin | 100 | 1×MIC | $H_2O$ |
| JH13 | Chloramphenicol | 62.5 | 5×MIC | DMSO |
| JH14 | Tigecycline | 6.25 | 5×MIC | DMSO |
| JH15 / 15* | Tetracycline† | 6.25 | 5×MIC | DMSO |
| JH16 | Azithromycin | 7.8 | 5×MIC | DMSO |
| JH17 | Gentamicin sulfate | 1.95 | 5×MIC | $H_2O$ |
| JH18 | Eravacycline | 0.2 | 5×MIC | $H_2O$ |
| JH20 | Sulfamonomethoxine | 1000 | 10×MIC | DMSO |
| JH21 | Sulfisoxazole | 1000 | 5×MIC | DMSO |
| JH23 | Trimethoprim | 6.25 | 5×MIC | DMSO |
| JH24 | Tetroxoprim | 62.5 | 5×MIC | DMSO |
| JH26 | Gatifloxacin | 0.31 | 5×MIC | DMSO |
| JH27 | Temafloxacin | 0.78 | 5×MIC | DMSO + HCl (16.6 mM) |
| JH28 | Lascufloxacin | 0.39 | 5×MIC | DMSO |
| JH29 | Ciprofloxacin | 0.16 | 5×MIC | $H_2O$ + HCl (0.1 M) |
| JH30 | Levofloxacin | 0.31 | 5×MIC | DMSO |
| JH31 / 31* | Nitrofurantoin† | 250 | 5×MIC | DMSO |
| JH32 | Gepotidacin | 25 | 10×MIC | DMSO |
| JH33 | Nalidixic acid | 250 | 5×MIC | DMSO |
| JH35 | Flumequine | 12.5 | 5×MIC | DMSO |
| JH36 | Enrofloxacin | 0.39 | 5×MIC | DMSO |
| JH37 | Rifaximin | 125 | 5×MIC | DMSO |
| JH38 | Rifampicin | 62.5 | 5×MIC | DMSO |
| JH39 | Rifabutin | 62.5 | 5×MIC | DMSO |
| JH40 | Triclosan | 15.5 | 50×MIC | DMSO |
| JH41 | Cerulenin | 200 | 1×MIC | DMSO |
| JH42 | MUT056399 | 15.6 | 10×MIC | DMSO |
| JH43 | Colistin | 4.6875 | 15×MIC | $H_2O$ |
| JH44 | Polymyxin B | 2.34 | 30×MIC | DMSO + HCl (16.6 mM) |
| JH45 | Squalamine | 3.12 | 1×MIC | DMSO |
| JH46 | Thanatin | 6.25 | 10×MIC | $H_2O$ |
| JH47 | Lolamicin | 12.5 | 5×MIC | DMSO |
| JH48 | PF-5081090 | 1.55 | 5×MIC | DMSO |
| JH49 | ACHN-975 | 0.78 | 5×MIC | DMSO |
| JH50 | CHIR-090 | 1.55 | 5×MIC | DMSO |
| JH51 | Xanthocillin | 1.56 | 1×MIC | DMSO |
| JH53 | Nitroxoline | 12.5 | 1×MIC | DMSO |
| JH55 | Kanamycin | 7.81 | 5×MIC | $H_2O$ |
| JH56 | Tobramycin | 7.81 | 5×MIC | $H_2O$ |
| JH58 / 58* | Nitrofurazone† | 500 | 20×MIC | DMSO |
| JH61 | Debio-1452-$NH_2$ | 93.75 | 15×MIC | DMSO |
| JH62 | BB-78485 | 15.63 | 10×MIC | DMSO |
| JH64 | D12 | 250 | 5×MIC | DMSO |
| JH65 / 65* | D8† | 125 | 10×MIC | DMSO |
| JH66 | Darobactin D22 | 39.06 | 50×MIC | DMSO |
| JH67 | Myxovirescin | 7.8 | 5×MIC | DMSO |
| JH68 | Cystobactamid CN-DM-861 | 1.56 | 5×MIC | DMSO |

Table S4: DIA-PASEF scan windows used for timsTOF Pro acquisition, including ion mobility range ($1/K_0$) and scan width (m/z).

| **MS Type** | **Scan** | **Start IM** [$1/K_0$] | **End IM** [$1/K_0$] | **Start Mass** [m/z] | **End Mass** [m/z] |
|---|---|---|---|---|---|
| MS1 | 0 | 0.60 | 1.60 | 100 | 1700 |
| dia-PASEF | 1 | 0.90 | 1.20 | 800 | 826 |
| dia-PASEF | 1 | 0.60 | 0.90 | 400 | 426 |
| dia-PASEF | 2 | 0.92 | 1.22 | 825 | 851 |
| dia-PASEF | 2 | 0.62 | 0.92 | 425 | 451 |
| dia-PASEF | 3 | 0.93 | 1.23 | 850 | 876 |
| dia-PASEF | 3 | 0.63 | 0.93 | 450 | 476 |
| dia-PASEF | 4 | 0.95 | 1.25 | 875 | 901 |
| dia-PASEF | 4 | 0.65 | 0.95 | 475 | 501 |
| dia-PASEF | 5 | 0.96 | 1.26 | 900 | 926 |
| dia-PASEF | 5 | 0.66 | 0.96 | 500 | 526 |
| dia-PASEF | 6 | 0.98 | 1.28 | 925 | 951 |
| dia-PASEF | 6 | 0.68 | 0.98 | 525 | 551 |
| dia-PASEF | 7 | 0.99 | 1.29 | 950 | 976 |
| dia-PASEF | 7 | 0.69 | 0.99 | 550 | 576 |
| dia-PASEF | 8 | 1.01 | 1.31 | 975 | 1001 |
| dia-PASEF | 8 | 0.71 | 1.01 | 575 | 601 |
| dia-PASEF | 9 | 1.02 | 1.32 | 1000 | 1026 |
| dia-PASEF | 9 | 0.72 | 1.02 | 600 | 626 |
| dia-PASEF | 10 | 1.04 | 1.34 | 1025 | 1051 |
| dia-PASEF | 10 | 0.74 | 1.04 | 625 | 651 |
| dia-PASEF | 11 | 1.06 | 1.36 | 1050 | 1076 |
| dia-PASEF | 11 | 0.76 | 1.06 | 650 | 676 |
| dia-PASEF | 12 | 1.07 | 1.37 | 1075 | 1101 |
| dia-PASEF | 12 | 0.77 | 1.07 | 675 | 701 |
| dia-PASEF | 13 | 1.09 | 1.39 | 1100 | 1126 |
| dia-PASEF | 13 | 0.79 | 1.09 | 700 | 726 |
| dia-PASEF | 14 | 1.10 | 1.40 | 1125 | 1151 |
| dia-PASEF | 14 | 0.80 | 1.10 | 725 | 751 |
| dia-PASEF | 15 | 1.12 | 1.42 | 1150 | 1176 |
| dia-PASEF | 15 | 0.82 | 1.12 | 750 | 776 |
| dia-PASEF | 16 | 1.13 | 1.43 | 1175 | 1201 |
| dia-PASEF | 16 | 0.83 | 1.13 | 775 | 801 |

Table S5: Composition of buffers and media referenced in the main Methods but not defined inline.

| **Buffer/Medium** | **Composition** |
|---|---|
| PBS | 10 mM $Na_2HPO_4$, 1.8 mM $KH_2PO_4$, 140 mM NaCl, 2.7 mM KCl in water, pH 7.4 |
| LB medium (Carl Roth) | 0.5% (w/v) yeast extract, 1.0% (w/v) peptone, 0.5% (w/v) NaCl in water, pH 7.5 |

Table S6: Text vs. No-text statistical analysis

| Model | Metric | Δ mean | 95% CI low | 95% CI high | $p_{\text{Bonf}}$ |
|---|---|---|---|---|---|
| LightGBM | macro-F1 | 0.1253 | 0.0710 | 0.1797 | $2.29 \times 10^{-4}$ |
| LightGBM | macro PR AUC OvR | 0.1065 | 0.0765 | 0.1366 | $4.40 \times 10^{-7}$ |
| LightGBM | macro ROC AUC OvR | 0.0384 | 0.0254 | 0.0514 | $8.21 \times 10^{-6}$ |
| PyTorch-MLP | macro-F1 | 0.1779 | 0.1309 | 0.2250 | $1.44 \times 10^{-7}$ |
| PyTorch-MLP | macro PR AUC OvR | 0.1125 | 0.0789 | 0.1461 | $1.13 \times 10^{-6}$ |
| PyTorch-MLP | macro ROC AUC OvR | 0.0417 | 0.0239 | 0.0596 | $1.96 \times 10^{-4}$ |

Table S7: Top 3 model–feature combinations for all models, reported with mean PR AUC and 95% confidence intervals across cross-validation cycles.

| Model | Coalition | Mean PR AUC | 95% CI |
|---|---|---|---|
| PyTorch-MLP | CellPaint + ECFP + Proteomics | 0.9519 | [0.9319, 0.9720] |
| PyTorch-MLP | CellPaint + ECFP + GC + MIC + Proteomics | 0.9477 | [0.9276, 0.9679] |
| PyTorch-MLP | CellPaint + ECFP + MIC + Proteomics | 0.9441 | [0.9248, 0.9634] |
| TabM | CellPaint + ECFP + Proteomics | 0.9575 | [0.9359, 0.9790] |
| TabM | CellPaint + ECFP + GC + MIC + Proteomics | 0.9569 | [0.9347, 0.9792] |
| TabM | CellPaint + ECFP + GC + Proteomics | 0.9568 | [0.9349, 0.9787] |
| LightGBM | CellPaint + ECFP + GC + MIC + Proteomics | 0.9525 | [0.9383, 0.9668] |
| LightGBM | CellPaint + GC + MIC + Proteomics | 0.9498 | [0.9332, 0.9663] |
| LightGBM | CellPaint + ECFP + Proteomics | 0.9478 | [0.9302, 0.9655] |
| MLP | CellPaint + ECFP + Proteomics | 0.9205 | [0.8997, 0.9414] |
| MLP | CellPaint + ECFP + GC | 0.9203 | [0.9007, 0.9400] |
| MLP | CellPaint + ECFP + GC + MIC + Proteomics | 0.9184 | [0.8955, 0.9413] |
| RF | CellPaint + ECFP + GC + MIC + Proteomics | 0.9243 | [0.8994, 0.9491] |
| RF | CellPaint + ECFP + GC + Proteomics | 0.9240 | [0.8994, 0.9485] |
| RF | CellPaint + ECFP + MIC + Proteomics | 0.9230 | [0.8967, 0.9493] |
| XGBoost | CellPaint + ECFP + Proteomics | 0.9396 | [0.9272, 0.9519] |
| XGBoost | CellPaint + ECFP + MIC + Proteomics | 0.9359 | [0.9167, 0.9551] |
| XGBoost | CellPaint + ECFP + GC + Proteomics | 0.9355 | [0.9177, 0.9533] |

Table S8: Figure 5 overall method performance (LOO/LOCO uncertainty benchmark).

| Method | Coverage | Precision | Error recall | F1 |
|---|---|---|---|---|
| Meta-model (LR+Poly) | 0.441 | 0.689 | 0.721 | 0.705 |
| Low max score | 0.431 | 0.636 | 0.651 | 0.644 |
| Z-Gate (class-conditional) | 0.196 | 0.550 | 0.256 | 0.349 |
| High predictive entropy | 0.176 | 0.667 | 0.279 | 0.393 |
| High expected entropy | 0.157 | 0.562 | 0.209 | 0.305 |
| Z-Gate (global) | 0.137 | 0.500 | 0.163 | 0.246 |

#### References

[1] OpenAI. ChatGPT. https://chatgpt.com/ (2025). Used August 8, 2025.

[2] Schuh, M. G., Boldini, D. & Sieber, S. A. Synergizing Chemical Structures and Bioassay Descriptions for Enhanced Molecular Property Prediction in Drug Discovery. *Journal of Chemical Information and Modeling* **64**, 4640–4650 (2024). URL https://doi.org/10.1021/acs.jcim.4c00765.

[3] Devlin, J., Chang, M.-W., Lee, K. & Toutanova, K. BERT: Pre-training of Deep Bidirectional Transformers for Language Understanding (2019). URL http://arxiv.org/abs/1810.04805. ArXiv:1810.04805 [cs].

[4] Lee, J. *et al.* BioBERT: a pre-trained biomedical language representation model for biomedical text mining. *Bioinformatics* **36**, 1234–1240 (2020). URL https://doi.org/10.1093/bioinformatics/btz682.

[5] Gorishniy, Y., Kotelnikov, A. & Babenko, A. TabM: Advancing Tabular Deep Learning with Parameter-Efficient Ensembling (2025). URL http://arxiv.org/abs/2410.24210. ArXiv:2410.24210 [cs].

[6] Paszke, A. *et al.* PyTorch: An Imperative Style, High-Performance Deep Learning Library (2019). URL http://arxiv.org/abs/1912.01703. ArXiv:1912.01703 [cs].

[7] Pedregosa, F. *et al.* Scikit-learn: Machine Learning in Python (2018). URL http://arxiv.org/abs/1201.0490. ArXiv:1201.0490 [cs].

[8] Ke, G. *et al.* Guyon, I. *et al.* (eds) *LightGBM: A Highly Efficient Gradient Boosting Decision Tree.* (eds Guyon, I. *et al.*) *Advances in Neural Information Processing Systems*, Vol. 30 (Curran Associates, Inc., 2017). URL https://proceedings.neurips.cc/paper_files/paper/2017/file/6449f44a102fde848669bdd9eb6b76fa-Paper.pdf.

[9] Chen, T. & Guestrin, C. Krishnapuram, B. *et al.* (eds) *Xgboost: A scalable tree boosting system.* (eds Krishnapuram, B. *et al.*) *Proceedings of the 22nd ACM SIGKDD International Conference on Knowledge Discovery and Data Mining*, KDD '16, 785–794 (Association for Computing Machinery, New York, NY, USA, 2016). URL https://doi.org/10.1145/2939672.2939785.

[10] Ash, J. R. *et al.* Practically Significant Method Comparison Protocols for Machine Learning in Small Molecule Drug Discovery. *Journal of Chemical Information and Modeling* **65**, 9398–9411 (2025). URL https://doi.org/10.1021/acs.jcim.5c01609.

[11] Köllen, M. F. *et al.* Generative Deep Learning Pipeline Yields Potent Gram-Negative Antibiotics. *JACS Au* **5**, 4249–4259 (2025). URL https://doi.org/10.1021/jacsau.5c00602.

[12] Parker, E. N. *et al.* Implementation of permeation rules leads to a FabI inhibitor with activity against Gram-negative pathogens. *Nature Microbiology* **5**, 67–75 (2020).

[13] Shao, D. *et al.* Design, Synthesis, and Cytotoxic Activity of 3-Aryl-N-hydroxy-2-(sulfonamido)propanamides in HepG2, HT-1080, KB, and MCF-7 Cells. *Chemistry & Biodiversity* **16**, e1800646 (2019). URL https://onlinelibrary.wiley.com/doi/abs/10.1002/cbdv.201800646. _eprint: https://onlinelibrary.wiley.com/doi/pdf/10.1002/cbdv.201800646.

[14] Hübner, I. *et al.* Broad Spectrum Antibiotic Xanthocillin X Effectively Kills *Acinetobacter baumannii via* Dysregulation of Heme Biosynthesis. *ACS Central Science* **7**, 488–498 (2021). URL https://pubs.acs.org/doi/10.1021/acscentsci.0c01621.
